# Endothelial Nucleoporin93 (Nup93) Maintains Vascular Function via Sun1-Dependent Regulation of RhoA-eNOS Signaling

**DOI:** 10.1101/2025.07.18.664980

**Authors:** Tung D. Nguyen, Yumna Z. Khan, Faruk Hossen, Riya Makim, Justin M. Banks, Julia Michalkiewicz, Michael A. Winek, Luis Henrique O. de Moraes, James C. Lee, Shane A. Phillips, Monica Y. Lee

## Abstract

As the innermost lining of blood vessels, endothelial cells (ECs) regulate blood flow, maintain vascular tone, and limit inflammation for vessel health. EC-derived nitric oxide (NO), synthesized by endothelial nitric oxide synthase (eNOS), is a vasodilator essential for improving blood flow and vascular homeostasis. The RhoA/ROCK pathway regulates eNOS levels, where overactivation decreases eNOS expression and downstream NO production. As such, RhoA/ROCK hyperactivity and increased pMLC have been identified as major contributors to age-associated vasoconstriction and hypertension. Intriguingly, recent studies identify Sun1, a key component of the linker of nucleoskeleton and cytoskeleton (LINC) complex, as a major regulator of RhoA/ROCK activity. Moreover, endothelial aging deteriorates nuclear pore complexes (NPCs) (*i.e.* nucleoporin [Nup93]) and impairs nucleocytoplasmic transport, thereby insinuating a role for nuclear envelope components in vessel homeostasis. Here, we show that targeted loss of endothelial Nup93 in adult mice significantly reduces eNOS expression and NO bioavailability for consequent defects in NO-dependent vasodilatory responses. *In vitro* knockdown of Nup93 in primary human ECs also decreases both eNOS expression and NO production. Mechanistically, we find that loss of Nup93 significantly reduces endothelial Sun1 levels for a concomitant increase in RhoA activity. Indeed, restoring Sun1 protein levels in Nup93-deficient ECs mitigates RhoA activity to rescue both eNOS expression and NO production. Taken together, we demonstrate endothelial Nup93, through Sun1 stabilization, as a novel regulator of eNOS-NO signaling and vessel reactivity, contributing to the growing importance of nuclear membrane components in EC and vascular biology.

## INTRODUCTION

Endothelial cells (ECs) serve as a dynamic and selective barrier between tissue and circulating blood components. Endothelial activation, an initiating event for cardiovascular disease (CVD), promotes disease progression by triggering EC inflammation, vascular permeability, and consequent atherosclerotic lesion formation. Research in the last several decades unequivocally establishes endothelial nitric oxide synthase (eNOS) activation, and subsequent nitric oxide (NO) production, as key events in safeguarding against the development of CVD. As such, eNOS impairment, characteristic of human aging, diminishes NO bioavailability to trigger multiple vascular complications^1,2^. While maintaining eNOS function is well-known to mitigate CVD development, it remains entirely unknown how endothelial nuclear integrity, a key safeguard for EC health, influences eNOS function.

The nuclear envelope has garnered rising interest as a novel regulator of EC health and function. Within the nuclear envelope spans the linker of nucleoskeleton and cytoskeleton (LINC) complex, a structure that connects the cytoskeletal network to the nucleus to create a mechanically coupled unit^3–5^. Recent studies identify Sun1, a component of the LINC complex residing at the inner nuclear membrane, as an unconventional regulator of EC junctional stability and barrier function. Mechanistically, loss of endothelial Sun1 was shown to indirectly augment RhoA/ROCK activity through microtubule-based regulation of RhoGEF-H1, thereby identifying Sun1 as a suppressor of RhoA/ROCK signaling^6^. These studies highlight Sun1 in the regulation of peripheral EC events to identify nuclear envelope components as mediators of long-range cellular communication. Emerging studies have also implicated the nuclear pore complex (NPC), one of the largest structures of the nuclear envelope, and its component nucleoporin proteins in cardiovascular function^7,8^. As the major gateway for bidirectional nucleocytoplasmic exchange, NPCs permit the rapid diffusion of small biomolecules (<40kDa), whereas larger molecules require an active carrier-based transport system (*e.g.,* importins, exportins)^9–11^. Necessary for the NPC structure, nucleoporin93 (Nup93) is one of the most abundant nucleoporins in mammalian cells known to decline with age^12,13^. We recently provide striking evidence for age-associated Nup93 loss in ECs, adding vascular cells to the growing list of cell types that exhibit nuclear permeability with age^12,14^. Although NPCs are considered one of the largest components of the nuclear envelope, the impact of nucleoporin dysregulation on blood vessel function and vascular disease has not been investigated.

Decades of investigative work have uncovered various mechanisms of eNOS regulation where alternative, unanticipated regulators of eNOS continue to be uncovered^15,16^. Mechanistically, RhoA hyperactivity is one of the few mechanisms known to diminish eNOS function both at the transcriptional and post-translational level; eNOS mRNA and activation (*i.e.,* phosphorylation at the S1177 activation site) are drastically reduced with elevated RhoA signaling^17,18^. As such, targeting RhoA/ROCK activity has been a conceptually appealing therapeutic strategy for hypertension and other CVDs. While RhoA/ROCK hyperactivity is fully recognized as a key effector of eNOS dysregulation and vascular inflammation, the mechanisms of RhoA/ROCK regulation via nuclear envelope components are only now beginning to be elucidated.

The present study sought to determine the relationship between suboptimal endothelial Nup93 levels and eNOS function. Using both *in vitro* and *in vivo* approaches, we find that Nup93 loss, a hallmark of endothelial aging, triggers a corresponding decline in Sun1 levels, thereby driving aberrant RhoA/ROCK hyperactivity. As a result, ECs exhibit increased cellular stiffening and eNOS deficiency, leading to barrier permeability, decreased NO production, and impaired vessel reactivity. We reveal that restoration of Sun1 levels in Nup93-deficient ECs attenuates stress fiber formation, thereby rescuing EC barrier integrity and eNOS function. We have therefore identified the presence of a regulatory mechanism between endothelial NPC components and eNOS function, demonstrating nuclear envelope dysregulation as an instigator of eNOS disruption to illuminate previously unrecognized effectors of NO bioavailability and vessel health.

## MATERIALS AND METHODS

### Data Availability

Data supporting the findings in this study are available from the corresponding author upon reasonable request.

### Animal Procedures

Nup93 floxed mice in the C57BL/6J background were generated by flanking exon 2 of Nup93 with loxP sites via CRISPR/Cas9 methods utilizing a single-strand DNA as the repair template for loxP insertion. Nup93 floxed mice were validated by NGS amplicon sequencing and further confirmed using outside, genomic-specific PCR primers (**Table S1**). Additional backcrosses to C57BL/6J mice were performed to clean up potential off-targets. Conditional EC-specific Nup93 deletion (Nup93ECKO) mice were generated by crossbreeding homozygous Nup93 floxed mice with Cdh5 promoter-driven inducible Cre recombinase (Cdh5-Cre-ERT2, *R. Adams*) mice^19,20^. Conditional young adults (∼4-5 weeks of age) were injected with tamoxifen (100µg/g body weight) for 5 consecutive days via intraperitoneal delivery and maintained on a standard laboratory diet (Teklad 7912, Envigo) as previously described^21^. Both male and female mice were used for the present studies. At harvest, mice were euthanized via prolonged isoflurane inhalation followed by exsanguination under anesthesia. All animal experiments were approved by the Institutional Animal Care and Use Committee (IACUC) at the University of Illinois at Chicago.

### Isolation of Mouse Lung Endothelial Cells (MLECs)

MLECs were isolated as previously described^22–24^. In brief, mice were anesthetized as described above and perfused with PBS (pH 7.4) via the left ventricle prior to removal of lung tissue. Perfused lungs were finely minced and digested in 300U/mL Type I Collagenase/PBS solution (37°C, 1hr). Digested tissues were passed through a 70µm cell strainer, pelleted, and resuspended in sterile 0.1% BSA/PBS solution. Magnetic Dynabeads (Invitrogen, 11035) were pre-conjugated with an anti-mouse PECAM-1 (BD553370) antibody and used for EC enrichment (room temp., 15min). Bound ECs were washed in PBS and harvested for protein analysis using a RIPA buffer-based lysis solution. Protein concentrations were then determined using a standard Bradford Colorimetric Assay kit (Bio-Rad, 5000116).

### Tissue Immunostaining

The immunofluorescence staining of mouse tissues was performed as previously described^25^. In brief, mice were anesthetized and perfused with PBS (pH 7.4) prior to removal of the target organs (*e.g.* heart). Samples were fixed with 10% Formalin/PBS solution (4°C, overnight) followed by paraffin processing and embedding. Tissues were sectioned (5µm), permeabilized with 0.2% Triton X-100/PBS (room temp., 10min), and in blocking solution (5% BSA/PBS, room temp., 1hr). Adjacent tissue sections were incubated with the following primary antibodies (4°C, overnight): anti-Nup93 (1:250, Santa Cruz, sc374399); anti-phospho-myosin light chain (T18/S19) (1:250, Cell Signaling, 3674S); anti-VWF (1:250, Abcam, ab11713). Tissue sections were then incubated with corresponding Alexa Fluor secondary antibodies (room temp., 2hrs), and stained with DAPI. Images were obtained using an inverted fluorescent microscope (Leica DMi8) and quantified using ImageJ software. Signal intensities were normalized relative to control conditions.

### Evan’s Blue Miles Assay

The Miles assay was used to assess baseline and stimuli-induced permeability, as previously published^26^. In brief, mice were subjected to intraperitoneal injection of pyrilamine maleate salt (40mg/kg body weight in 0.9% saline; Sigma-Aldrich) to inhibit the release of endogenous histamine. Mice were anesthetized and injected with sterile Evans blue (EB) dye (50mg/kg body weight) via retro-orbital delivery. After 10 minutes, local permeability was induced via intradermal bolus injections (50uL) of VEGF-A165 (PeproTech, 200ng) or histamine (CAT, 125ug) on shaved back skin. PBS was also injected (50uL) as controls. Mice were placed at 37C where EB dye was allowed to circulate for 30 minutes. Mice were subsequently harvested for removal of target organs (*e.g.* heart, lungs, liver, kidney, pancreas, inguinal fat, spleen, back skin). Samples were initially weighed (‘wet weight’), allowed to dry in a gravity convection oven (55°C, overnight), and weighed again (‘dry weight’) to evaluate tissue edema (% wet/dry weight) as a secondary readout of vascular leakage (REF). Dried tissues were then incubated in 100% formamide (55°C, overnight) for extraction of EB dye. The amount of EB in each tissue sample was quantified by measuring absorbance at 610nm against a standard curve. Tissue EB and edema values are presented relative to control conditions.

### Flow-Mediated Vessel Dilation

Mice were anesthetized, as described above, and perfused with PBS (pH 7.4) prior to removal of mesenteric arteries. Isolated vessels were carefully excised and placed in 1M HEPES solution on ice. Mesenteric arteries between 100-200µm were cannulated using glass micropipettes filled with Krebs buffer (pH 7.4) and equilibrated at 60 cmH_2_O in a temperature-controlled chamber (37°C, 30min). Arteries were visualized using an inverted brightfield microscope (Olympus CKX41) and the baseline diameter of selected arteries was recorded using an image measurement system (Boekeler model VIA-100). Endothelin-1 (120pM) was introduced to constrict the chosen arteries up to 50% of their baseline diameter, and flow-induced vasodilation was achieved by subsequently exposing the arteries to incremental pressure gradients of 20, 40, 60, and 100 cmH_2_O^27,28^. Arterial diameter was measured after 5 minutes under each pressure gradient to determine the % change in vessel diameter in response to increasing flow.

### Cell Culture

Primary human retinal endothelial cells (HRECs) were purchased (Cell Systems, ACBRI-181-V) and cultured in endothelial cell growth media (Lonza, CC-3162) supplemented with 5% FBS (Corning, 35-011-CV), 1% L-glutamine, and 1% penicillin-streptomycin (37°C, 5% CO_2_)^14^. Confluent HRECs were subcultured every 3 days, and cells before passage 10 were used for all *in vitro* experiments reported herein. Human embryonic kidney 293T (HEK293T) cells were utilized for lentiviral packaging and cultured in high glucose DMEM (Lonza, BE12-709F) supplemented with 10% FBS, 1% L-glutamine, and 1% penicillin-streptomycin.

### Lentiviral Transduction

Glycerol stocks of pLKO.1-puro lentiviral plasmid vectors containing shRNA targeting human Nup93 (NM_014669) and an empty vector insert (shEmpty) were purchased from Sigma-Aldrich. Lentivirus expressing shNup93 or shEmpty were generated by PEI-mediated co-transfection of pLKO.1-puro, psPAX2, and pMD2.G in HEK93T cells (72 hours), as previously described^14^. Human Nup93 (NM_014669), Sun1 (NM_025154), and an empty insert (Lenti–Empty vector) were subcloned in-house into a pCCL-PGK-MCS lentiviral plasmid vector (gift from Dr. Jan Kitajewski [UIC]). Both the human Nup93 and Sun1 lentiviral constructs were further designed to contain a FLAG tag at their respective C-terminus. Lentivirus expressing Nup93 (Nup93-FLAG), Sun1 (Sun1-FLAG), or Empty vector were generated as described. The Rev-Glucocorticoid Receptor-GFP (RGG) expression vector (gift from Dr. John Hanover^29^ [NIH]) was subcloned into a pLenti-puro lentiviral plasmid vector. Sub-confluent HRECs were transduced with the respective lentivirus (1:10 dilution in EC growth media, 20 hrs). HRECs infected by lentiviral vectors carrying puromycin-resistance were treated with 2.5µg/mL puromycin (Gibco, A11138-03) for 24 hours to eliminate non-transduced cells.

### siRNA Transfection

FlexiTube Premix solutions (20µM) containing siRNA targeting human Sun1 (NM_025154) or a negative control insert (siScramble) were purchased from Qiagen. Sub-confluent HRECs were transfected with the respective siRNA (20nM) via Oligofectamine Transfection Reagent (Invitrogen, 12252011) in reduced-serum Opti-MEM (Gibco, 31985070) for 6 hours (37°C, 5% CO_2_). siRNA-transfected HRECs were subsequently supplemented with EC growth media and 10% FBS for subsequent assays.

### Nitrate/Nitrite Assay

Mice were fasted overnight prior to retroorbital-based blood collection (∼200µL/mouse), as previously described^25^. In brief, blood samples were collected into Eppendorf tubes containing ∼4µL 0.5M EDTA (Fisher, BP120), gently mixed, and immediately placed on ice. Samples were centrifuged (1000xg, 4°C, 10min), where plasma supernatant fractions were collected for analyses. For *in vitro* measurements, confluent HRECs were incubated in phenol red-free growth media (Lonza, CC-3129) 24-hours prior to media collection. NO concentrations were determined using the Nitrate/Nitrite Colorimetric Assay Kit according to the manufacturer’s instructions (Caymen, 780001).

### Atomic Force Microscopy

Endothelial stiffness was assessed using an atomic force microscopy system (Asylum Research MFP-3D-BIO). In brief, a silicon nitride cantilever with a 5µm-diameter borosilicate glass particle tip (Novascan, PT.BORO.SN.5) was initially calibrated for contact mode force measurement. The cantilever (spring constant: 0.06N/m) was then configured to gradually descend towards individual cells and induce cell indentation (∼5-7nN), corresponding to ∼0.7-1.2µm indentation depth^30,31^. Cellular stiffness was determined in sub-confluent HRECs through fitting the force versus indentation curves at 15-20 distinct cell surface sites in 7-10 cells to the Hertz model to acquire the sample elastic modulus (kPa).

### Western Blotting

Cells were harvested for protein analysis using a RIPA-based lysis solution, and protein concentrations were determined using a standard Bradford Colorimetric Assay kit (Bio-Rad, 5000116) as previously described^14^. Proteins were separated by SDS-PAGE and transferred to 0.45µm nitrocellulose membranes. Membranes were blocked in 1% Casein/ TBS (room temp, 1hr) and incubated with the following primary antibodies (4°C, overnight): anti-Nup93 (1:1000, Sigma-Aldrich, HPA017937); anti-γH2AX (1:500, Sigma-Aldrich, 05-636); anti-H2AX (1:1000, Cell Signaling, 2595S); anti-phospho-myosin light chain (T18/S19) (1:500, Cell Signaling, 3674S); anti-myosin light chain (1:1000, Cell Signaling, 3672S); anti-PECAM-1 (1:1000, Santa Cruz, sc376764); anti-GAPDH (1:1000, Cell Signaling, 2118S); anti-phospho-eNOS (S1176) (1:500, BD Biosciences, 612392); anti-eNOS (1:1000, BD Biosciences, 610297); anti-β-Actin (1:5000, Sigma-Aldrich, A5441); anti-RhoA (1:1000, Santa Cruz, sc179); anti-Sun1 (1:1000, Invitrogen, MA5-34799); anti-Sun2 (1:1000, Invitrogen, PA5-51539). Corresponding DyLight secondary antibodies (Rockland) were introduced (room temp, 1hr) and visualized using the LI-COR Odyssey Imaging System. Using the ImageJ software, protein levels were normalized to housekeeping proteins, and normalized values are presented relative to controls.

### Immunofluorescence Staining

HRECs were fixed in 4% PFA/PBS (room temp., 10min) and permeabilized in 0.2% Triton X-100/PBS (room temp., 10min). Cells were then blocked in 5% BSA/PBS (room temp., 1hr) and incubated with the following primary antibodies (4°C, overnight): anti-Nup93 (1:750, Lusk Lab, Yale University); anti-Sun1 (1:250, Invitrogen, PA5-52564); anti-GFP (1:250, Santa Cruz, sc9996); anti-FLAG (1:250, Sigma-Aldrich, F1804). Cells were then incubated with corresponding Alexa Fluor secondary antibodies (room temp., 2hrs), and stained with DAPI. Images were visualized using an inverted fluorescent microscope (Leica DMi8) and quantified using the ImageJ software. To quantify nuclear and cytoplasmic localization, signal intensities in cell nuclei were calculated relative to DAPI staining. Cytoplasmic localization was quantified by subtracting nuclear signal from whole image intensity, and nuclear signals were then divided by cytoplasmic intensity in each condition for ratiometric presentation. Fluorescent intensities were normalized relative to control conditions.

### Gelatin Trapping Assay

*In vitro* permeability assays were performed, as previously described^32,33^. In brief, gelatin (Sigma-Aldrich, G1890) was dissolved in 0.1M NaHCO_3_ buffer (pH 8.3) to a final concentration of 10mg/mL. EZ-Link NHS-LC-LC-Biotin (Thermo Fisher, 21336) in sterile DMSO (5.7mg/mL) was conjugated to the gelatin/NaHCO_3_ solution to a final concentration of 0.25mg/mL, sterile-filtered through 0.22µm PES membrane, and stored at-20°C. Chambered slides (ibidi, 80821) were coated with biotinylated gelatin (4°C, overnight) followed by sterile PBS wash and primary HREC seeding. Streptavidin-Alexa Fluor 568 (1:500, Invitrogen, S11226) was added briefly to confluent HRECs (room temp., 30sec) prior to fixing, permeabilizing, and blocking as described above. Cells were incubated with anti-VE-cadherin (1:500, Invitrogen, 14-1449-82) overnight at 4°C followed by the introduction of corresponding Alexa Fluor secondary antibody and Phalloidin-Atto 488 (Sigma-Aldrich, 49409) to stain the actin filaments (room temp., 2hrs). Images were taken using an inverted fluorescent microscope (Leica DMi8) and quantified using the ImageJ software. Fluorescent intensities were normalized relative to controls.

### RhoA Activation Assay

*In vitro* RhoA activity was determined by measuring the level of GTP-bound RhoA in cultured cells according to the manufacturer’s instructions (Cytoskeleton, BK124). In brief, confluent HRECs were harvested for protein analysis using a RIPA-based lysis solution, and protein concentrations were determined using a standard Bradford Colorimetric Assay kit (Bio-Rad, 5000116). Proteins were added to RhoA-GTP affinity wells and incubated with anti-RhoA antibody (1:250, room temp., 45min). Proteins were then incubated with corresponding HRP-labeled secondary antibody (room temp., 45min) followed by introduction of HRP detection reagents (37°C, 15min). The level of GTP-bound RhoA in HRECs was assessed by measuring absorbance at 490nm and presented relative to control conditions.

### β-Galactosidase Senescence Assay

Senescence-associated β-Galactosidase (SA-βGal) was measured according to the manufacturer’s instructions (Cell Signaling, 9860S) and as previously described^14^. In brief, confluent HRECs were fixed with the provided 1X Fixative solution, washed in PBS, and incubated in β-Gal Staining Solution (pH 6.0) overnight at 37°C to develop the cyan color in senescent cells. Images were visualized using an inverted fluorescent microscope (Leica DMi8) and quantified using the ImageJ software.

### RNA Extraction and RT-qPCR

Isolation of human RNA samples and quantitative real-time PCR were performed as previously described^14^. In brief, confluent HRECs were harvested for total RNA using the RNeasy Mini Kit (Qiagen, 74104). RNA purity and concentration were assessed with a NanoDrop One instrument (Invitrogen). Total RNA (0.5µg) was converted into cDNA using the High-Capacity cDNA Reverse Transcription Kit (Applied Biosystems, 4368814). RT-qPCR was performed using the Fast SYBR Green Master Mix (Applied Biosystems, 4385612) using a ViiA 7 Real-Time PCR System (Applied Biosystems). Relative mRNA levels were normalized to GAPDH, and normalized levels were further expressed relative to controls. Primer sequences for human genes used herein are listed in **Table S2**.

### Chemicals and Reagents

Tumor necrosis factor alpha (TNFα) was purchased (Sigma, H8916) and dissolved in sterile Milli-Q water to a stock concentration of 10µg/mL for *in vitro* experiments. Y-27632 was purchased (Sigma, 688000) and dissolved in sterile Milli-Q water to a stock concentration of 10mM for *in vitro* experiments. S-nitroso-N-acetylpenicillamine (SNAP) was purchased (Sigma, N3398) and dissolved in sterile DMSO to a stock concentration of 0.2M for *in vitro* experiments. Tamoxifen was purchased (Sigma, T5648) and homogenized in sterile peanut oil to a stock concentration of 20mg/mL for *in vivo* experiments.

### Statistical Analysis

All data are presented as mean ± SEM and analyzed using GraphPad Prism v9.0 software. Statistical analyses reflect an unpaired two-tailed Student t-test for two-group comparisons of normally distributed data with equal variance. Multi-group comparisons with equal variances are performed using 2-way ANOVA followed by Tukey post-hoc test. A p-value<0.05 is considered statistically significant.

## RESULTS

### Endothelial Nup93 is necessary for barrier integrity and vascular tone regulation

Endothelial senescence is associated with pathological conditions, including increased vascular permeability and impaired vasodilation^34,35^. Our recent reports identify loss of Nup93 as an age-associated event in vascular ECs to suggest that endothelial NPC deterioration may precipitate adverse vascular events. We therefore generated an inducible EC-targeted Nup93 conditional mouse model to determine the consequences of suboptimal endothelial Nup93 in vascular homeostasis and function. In brief, Nup93 floxed mice were bred to the inducible Cdh5-CRE driver (Cdh5-Cre-ERT2)^19,20^ to generate an endothelial-targeted Nup93 murine system. Endothelial-targeted Nup93 depletion was achieved following a previously established tamoxifen regimen, where young adult mice (∼4wks of age) are administered tamoxifen for 5 consecutive days via intraperitoneal injection. Intriguingly, the targeted deletion of endothelial Nup93 in the established vasculature is incompatible with life, as Nup93iECKO mice exhibit lethality within 5 weeks of the last tamoxifen injection (**Fig.S1**). The observed lethality occurs irrespective of sex, thereby demonstrating the importance of endothelial Nup93 in the established vasculature. To investigate the vascular consequences of endothelial Nup93 loss, all adult studies herein were performed within 3 weeks of the last tamoxifen injection to avoid the mortality time window. Isolation of primary mouse lung ECs (MLECs), as confirmed through PECAM1 detection, indicates a significant loss of Nup93 expression at 3 weeks post-tamoxifen to reveal efficient Nup93 depletion in ECs using our inducible strategy (**Fig.1A&B**). In our recent reports, *in vitro* loss of Nup93 triggered cellular senescence and EC dysfunction^14^. Paralleling *in vitro* observations^14^, Nup93iECKO MLECs also exhibit reduced expression of LaminB1 and significantly elevated levels of γH2AX (*i.e.,* H2AX phosphorylated at Ser139), both well-established markers of cell senescence (**Fig.1A-D**). We find that loss of Nup93 also increases phosphorylation (Thr18/Ser19) levels of MLC (**Fig.1E**), a downstream effector of RhoA/ROCK activity^36^. Cross-sectional analysis of the coronary vasculature was also performed, where immunofluorescent-based detection of endothelial Nup93, as delineated by VWF expression, is significantly reduced in Nup93iECKO mice when compared to WT controls, further validating our deletion strategy (**Fig.1F-H**). Similar to our immunoblot analysis, Nup93iECKO mice exhibit increased MLC phosphorylation in the endothelial layer (*arrows*). MLC phosphorylation was also markedly increased in vascular regions beyond the endothelium to suggest that other cell types involved in mediating vessel tone (*e.g.,* vascular smooth muscle cells) may also be adversely impacted with endothelial Nup93 loss.

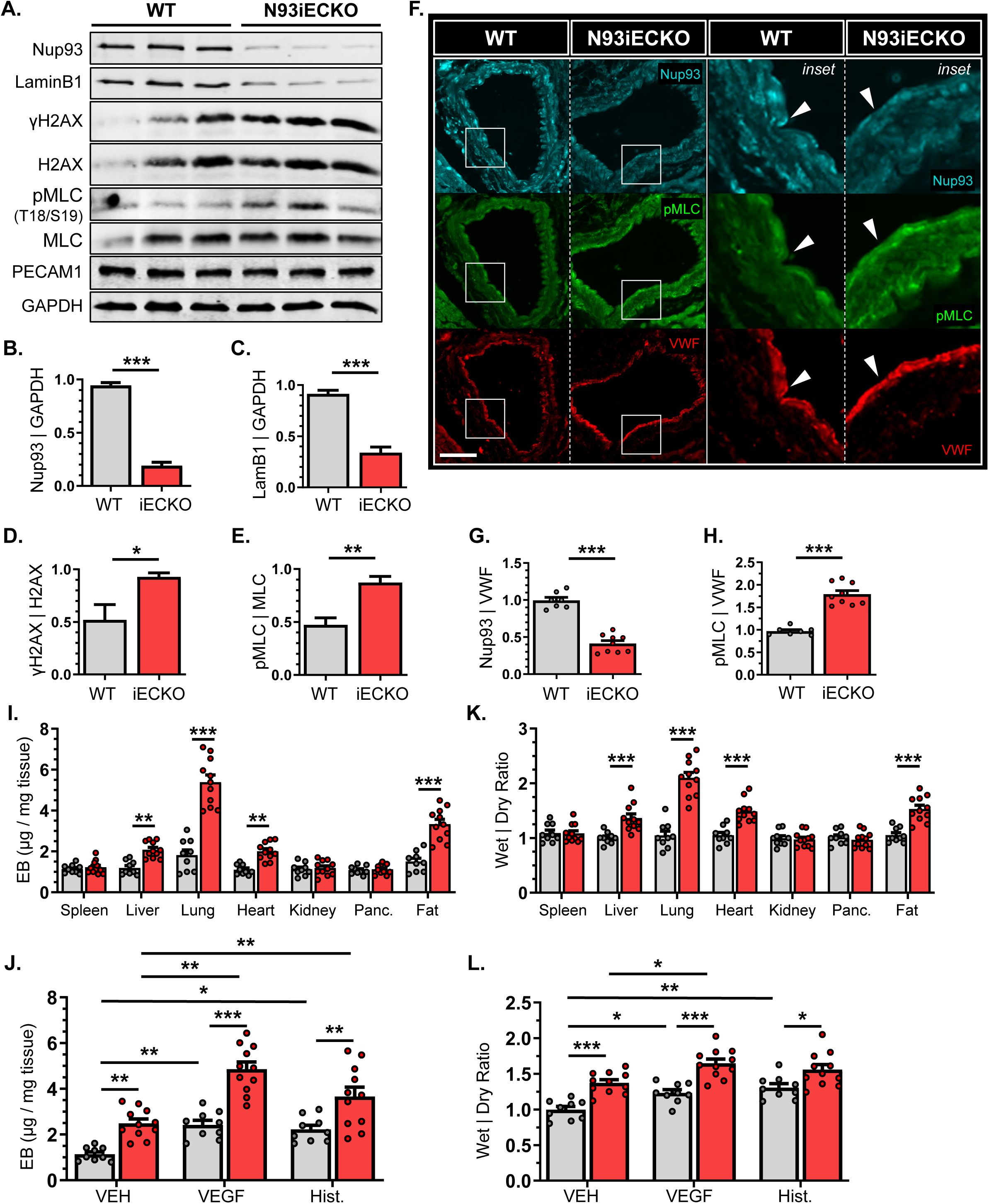

Activation of endothelial RhoA/ROCK/MLC signaling has been previously shown to alter the cellular cytoskeleton, disrupting cell-cell adhesion for consequent vessel permeability. To examine the consequence of endothelial Nup93 loss on barrier integrity, mice (Nup93iECKO and WT controls) were intravascularly injected with Evans Blue (EB) dye, whereupon a panel of organs were isolated to measure EB tissue extravasation as a readout of vascular leakage^26^. Compared to WT controls, Nup93iECKO mice exhibited a significant increase in EB dye extravasation across several highly vascularized tissues, namely the lungs, heart, liver, and inguinal fat to indicate increased baseline permeability (**Fig.1I**). Mice were also subjected to intradermal injections of VEGF-A (4µg/mL, 200ng) or histamine (2.5mg/mL, 125ng) to measure local skin permeability in response to factors. Both VEGF and histamine rely on endothelial RhoA/ROCK signal activation, and actomyosin contraction, to elicit increases in vessel permeability^37–40^. Intradermal injections of either VEGF-A or histamine increased skin permeability in WT control mice, as expected. Agonist-induced permeability is further increased with loss of endothelial Nup93, as Nup93iECKO mice exhibit heightened EB dye levels in measured skin samples (**Fig.1J**). These differences in vessel permeability were also revealed when assessing the wet-to-dry weight ratios of isolated tissues. Calculated weight ratios were found to be significantly higher in vascularized organs (*e.g.,* lungs, heart, liver, inguinal fat) from Nup93iECKO mice (**Fig.1K**). Moreover, the back skin of Nup93iECKO mice yielded an increase in wet-to-dry ratios with all introduced agonists, including the PBS control (**Fig.1L**). Taken together, endothelial loss of Nup93 leads to vascular leakage and an abnormal accumulation of interstitial fluid in highly perfused organs.

The RhoA/ROCK pathway is largely implicated in the regulation of vessel tone and peripheral vascular resistance. Aberrant RhoA/ROCK activation has been shown to increase vessel stiffening and clinical hypertension. Mechanistically, increased endothelial RhoA activity decreases eNOS phosphorylation and expression, thereby blunting NO production^17,18^. Intriguingly, primary MLECs isolated from Nup93iECKO mice exhibit a significant decrease in both phosphorylated (Ser1176) and total eNOS levels (**Fig.2A-C**). Plasma samples from Nup93iECKO mice also indicate a significant reduction in circulating NO levels, as determined via total nitrite and nitrate^41–43^, when compared to WT controls (**Fig.2D**). More importantly, flow-induced vasodilatory responses, a phenomenon reliant on endothelial eNOS function, is severely impaired in isolated mesenteric arteries of Nup93iECKO when compared to WT controls; Nup93iECKO vessels are unable to dilate in response to high intraluminal pressures (**Fig.2E**). Taken together, these findings establish, for the first time, endothelial nucleoporins, and specifically Nup93, as a necessary element of vessel homeostasis.

**Figure 2.**
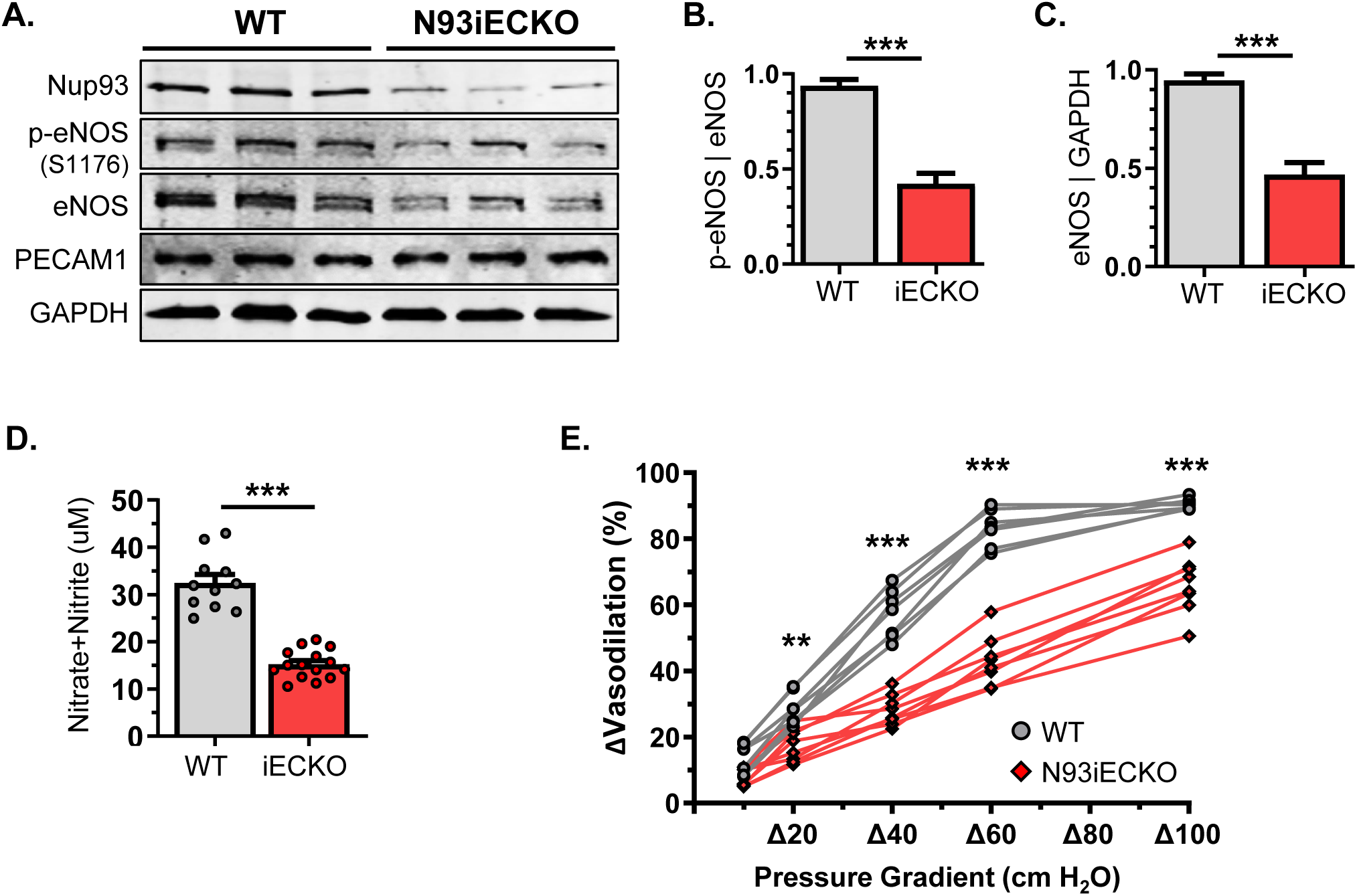
Endothelial Nup93 loss reduces eNOS expression and impairs vasodilatory function. **(A)** Immunoblotting of primary MLECs indicates a significant reduction in both phospho-eNOS (S1176) and total eNOS levels. (n=3-4 mice per lane, *shown in triplicates*). Quantified in **(B)** and **(C)**. **(D)** Nup93iECKO mice exhibit a corresponding decline in plasma NO levels. **(E)** Endothelial loss of Nup93 leads to a significant impairment in flow-induced dilation in isolated mesenteric arteries. (n=7 WT, n=8 Nup93iECKO). *\*\*\* p<0.001, ** p<0.01*.

### Loss of Nup93 promotes endothelial permeability and impairs eNOS function

Our *in vivo* findings identify a critical role for endothelial Nup93 in barrier stability. We therefore introduced a biotinylated gelatin assay for *in vitro*-based assessment of endothelial monolayer permeability, as previously described^32,44^. In brief, primary human retinal ECs were plated onto a biotinylated gelatin coating followed by selective knockdown of Nup93 using shRNA methods^14^. Endothelial loss of Nup93 leads to a significant increase in endothelial permeability, as Nup93-deficient ECs exhibit a significant increase in streptavidin signal (**Fig.3A&B**). VE-cadherin intensity at cellular junctions is also markedly reduced upon loss of Nup93, where similar decreases in expression are seen with other junctional proteins (**Fig.3A&C, Fig.S2**). Moreover, loss of Nup93 significantly increased stress fiber formation, a common outcome of RhoA hyperactivity (**Fig.3A&D**). Nup93 loss also decreases integrin β1 and increases total FAK protein levels (**Fig.S2D-G**), where recent studies implicate FAK in cellular tension transmission to the nucleus^45^. As such, we find that Nup93 knockdown leads to increased endothelial cellular stiffness, as assessed through AFM-based measurements (**Fig.3E**). These observations demonstrate Nup93 as a novel regulator of endothelial cytoskeletal architecture where loss of Nup93 leads to consequential EC rigidity.

**Figure 3.**
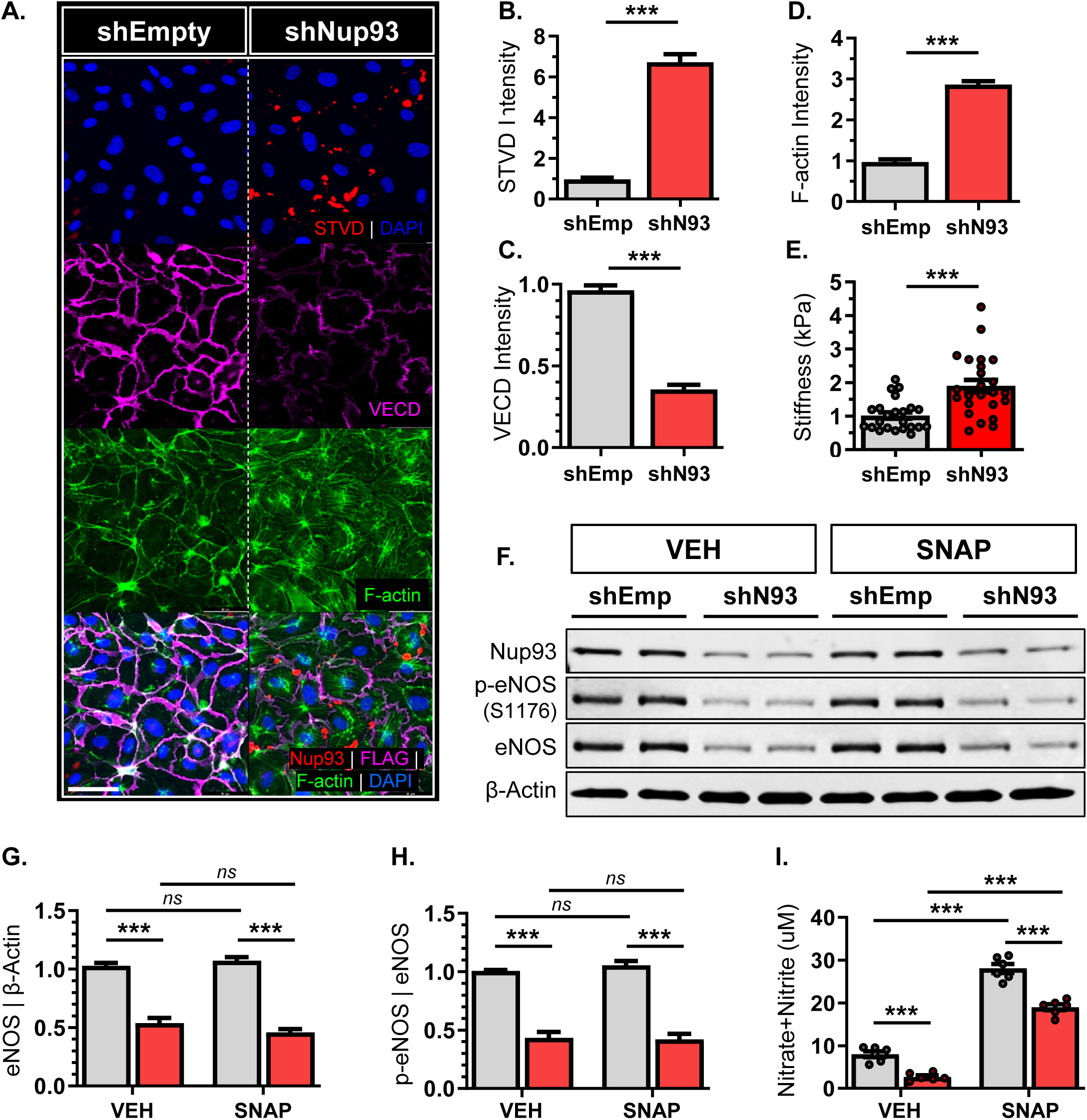
Nup93 knockdown compromises endothelial barrier function and impairs eNOS-derived NO production. Primary human ECs (shEmpty & shNup93) were plated on biotinylated gelatin. Immunofluorescence staining shows **(A)** increased paracellular permeability with Nup93 loss, as indicated by the visible streptavidin signal. Quantified in **(B)**. Nup93 knockdown ECs also exhibit **(C)** decreased VE-cadherin junctional localization and **(D)** increased stress fiber formation. **(E)** AFM measurements indicate a significant increase in endothelial stiffness with loss of Nup93. (n=3, 8 reads per group/expt). **(F)** Loss of Nup93 leads to a significant reduction in protein levels of both phospho-eNOS (S1176) and total eNOS, irrespective of SNAP (0.5μM, 24hrs) treatment *(shown in duplicates).* Quantified in **(G)** and **(H)**. **(I)** Conditioned media from Nup93-deficient ECs show a substantial decline in secreted NO levels. Scale bars=50µm. n=3, *\*\*\* p<0.001*

Age-associated arterial stiffening impairs the mechanical properties of the endothelium, thereby disrupting eNOS function for consequent reduction in nitric oxide production^46^. We therefore examined the impact of Nup93 loss on endothelial eNOS expression and function. In brief, ECs (shEmpty & shNup93) were treated with S-Nitroso-N-acetyl-DL-penicillamine (SNAP, 0.5μM, 1hr), an exogenous NO donor, followed by collection of both whole cell lysate and conditioned media. Endothelial Nup93 knockdown significantly decreased total eNOS protein expression as well as levels of the activating phosphorylation residue (S1176) (**Fig.3F-H**). SNAP treatment in both EC groups (shEmpty & shNup93) increased NO release, as expected (**Fig.3I**), with little effect on total eNOS protein. More importantly, Nup93-deficient ECs consistently exhibited decreased NO release to parallel the striking reduction in total eNOS protein expression (**Fig.3F&I**). These *in vitro* results parallel our *in vivo* observations to indicate a role for Nup93 in endothelial barrier function and eNOS-dependent NO production.

### Exogenous Nup93 protein expression levels re-establishes endothelial barrier integrity and eNOS function

The deterioration of Nup93 protein is strongly associated with cellular senescence across various cell types, including vascular ECs^14,47^. Establishing the importance of Nup93 protein levels, we recently demonstrated that restoring Nup93 protein expression in already senescent ECs was sufficient for endothelial reprograming from an inflammatory phenotype toward healthy ECs^14^. We therefore surmised that re-introducing Nup93 protein levels would also restore senescence-associated cytoskeletal defects. To model endothelial senescence *in vitro*, primary human ECs were initially treated with the inflammatory cytokine tumor necrosis factor alpha (TNFα) for 6 consecutive days^14,48^. As expected, chronic exposure to TNFα leads to a visible increase in streptavidin signal at intercellular regions to indicate paracellular permeability (**Fig.4A&B**). Long-term TNFα also induces stress fiber formation with concomitant decreases in VE-cadherin expression at cell-cell junctions (**Fig.4A,C,D**), consistent with previous reports detailing EC barrier dysfunction and senescence^49^. Similar to our recent reports^14^, lentiviral-based methods were then used for exogenous expression and restoration of Nup93 (**Fig.4E**). In brief, ECs were initially exposed to chronic inflammation (TNFα [10ng/mL]; 6 days) followed by lentiviral infection for exogenous Nup93 expression. These lentiviral methods achieve mild overexpression (∼1.3-fold), demonstrating the utility of the expression construct. Indeed, TNFα-exposed ECs show almost complete restoration of Nup93 protein to that of healthy controls (**Fig.4F&G**). More importantly, restoring Nup93 levels in senescent ECs significantly reduces streptavidin intensity while re-establishing both F-actin and VE-cadherin levels comparable to that of healthy controls (**Fig.4A-D**).

**Figure 4.**
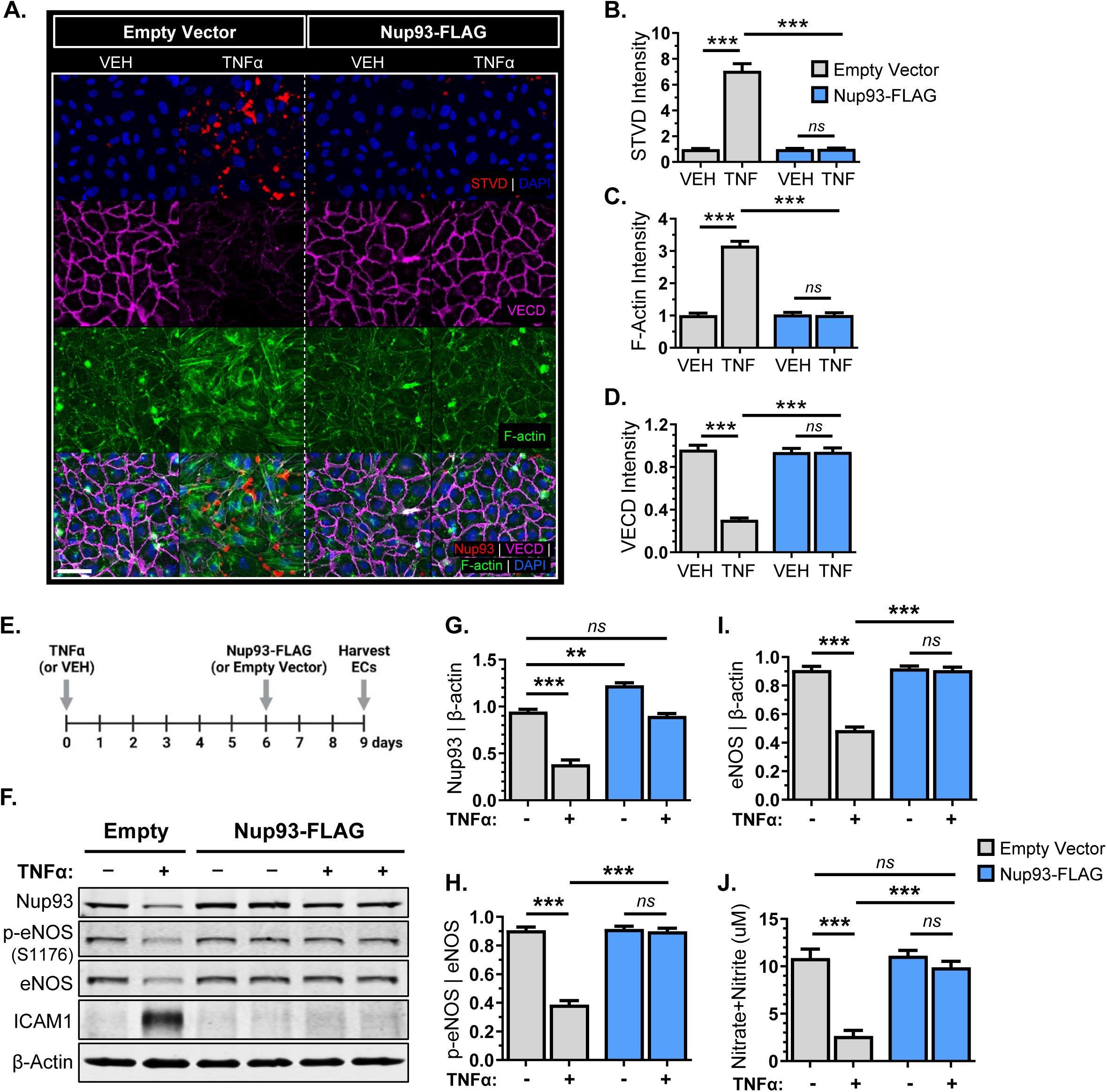
Restoring Nup93 levels re-establishes endothelial barrier integrity and eNOS function. **(A)** Chronic TNFα treatment (10ng/mL, 6 days) disrupts endothelial barrier function, as indicated by the visible paracellular streptavidin signal, decreased junctional VE-cadherin expression, and increased stress fibers. Quantified in **(B-D)**. **(E)** *Schematic*: HRECs were first exposed to the chronic inflammation model (TNFα [10 ng/mL]; 6 days) followed by lentiviral transduction for exogenous expression and restoration of Nup93 protein. **(F)** Chronic inflammation followed by exogenous Nup93 expression restores both eNOS phosphorylation and total protein levels. Quantified in **(G-I)**. **(J)** Chronic inflammation significantly decreases NO production, with rescue upon Nup93 restoration. Scale bars=50µm. n=3, *\*\*\* p<0.001, ** p<0.01*

Endothelial senescence is well-associated with a decrease in eNOS expression and function^2,50^. We therefore sought to determine whether exogenous expression of Nup93 could similarly restore eNOS function in senescent ECs. Using the same treatment strategy, ECs were harvested for both whole-cell lysate and conditioned media to assess eNOS protein and NO levels, respectively. Chronic inflammation significantly reduced endothelial Nup93 protein levels, similar to our previous reports^14^. Long-term TNFα treatment also decreased eNOS phosphorylation and total protein levels (**Fig.4F**), confirming previous reports of eNOS loss with chronic inflammation. More importantly, exogenous expression of Nup93 in already senescent ECs led to the recovery of both eNOS protein and baseline eNOS phosphorylation; phospho-eNOS and total eNOS match the levels of healthy control ECs (**Fig.4F,H,I**). Furthermore, restoring Nup93 protein levels in TNFα-exposed ECs led to full recovery of secreted NO similar to that of healthy ECs to indicate rescue in both eNOS protein and function (**Fig.4J**). These studies support the importance of endothelial Nup93, where suboptimal levels as a result of chronic inflammation triggers eNOS disruption via multiple mechanisms.

### Endothelial Nup93 deficiency leads to RhoA/ROCK hyperactivity

Stress fiber formation and cellular stiffening is largely associated with increased RhoA/ROCK signal activity^38,51^. The RhoA/ROCK pathway is also known to influence eNOS levels, where overactivation decreases eNOS expression and downstream NO production^17,18^. As such, RhoA/ROCK hyperactivity and increased substrate phosphorylation (*e.g*., MLC) have been identified as major contributors to vasoconstriction, clinical hypertension, and vascular aging. Given the known link between hyperactive RhoA signaling and stress fiber formation, we postulated a role for enhanced RhoA activity as a major contributor of endothelial barrier disruption in Nup93-deficient ECs. Here, we find that loss of endothelial Nup93 significantly elevates both RhoA protein levels and activity, as assessed through immunoblotting and enzyme immunoassays for GTP-bound RhoA, respectively (**Fig.5A-C**). Nup93-deficient ECs were hence treated with a pharmacological ROCK inhibitor (Y-27632), effectively blocking the downstream effector of RhoA signaling^52,53^. Consistent with our previous observations (**Fig.3**), endothelial knockdown of Nup93 increases paracellular leakage, decreases VE-cadherin expression, and enhances stress fiber formation (**Fig.5D-G**). Treatment with a pharmacological Rho/ROCK inhibitor (Y-27632, 10µM, 24hrs), however, reversed these effects; Y-27632 mitigates streptavidin signal and restores VE-cadherin junctional expression in Nup93-deficient ECs, similar to healthy EC controls (**Fig.5D-G**). F-actin intensity is also reduced in Nup93-deficient ECs upon Y-27362 exposure (**Fig.5D,G**), where AFM-based studies indicate a corresponding decrease in EC stiffness (**Fig.5H).** These findings implicate RhoA/ROCK hyperactivity as a consequence of endothelial Nup93 loss driving the detrimental impacts on endothelial barrier function and cellular rigidity.

**Figure 5.**
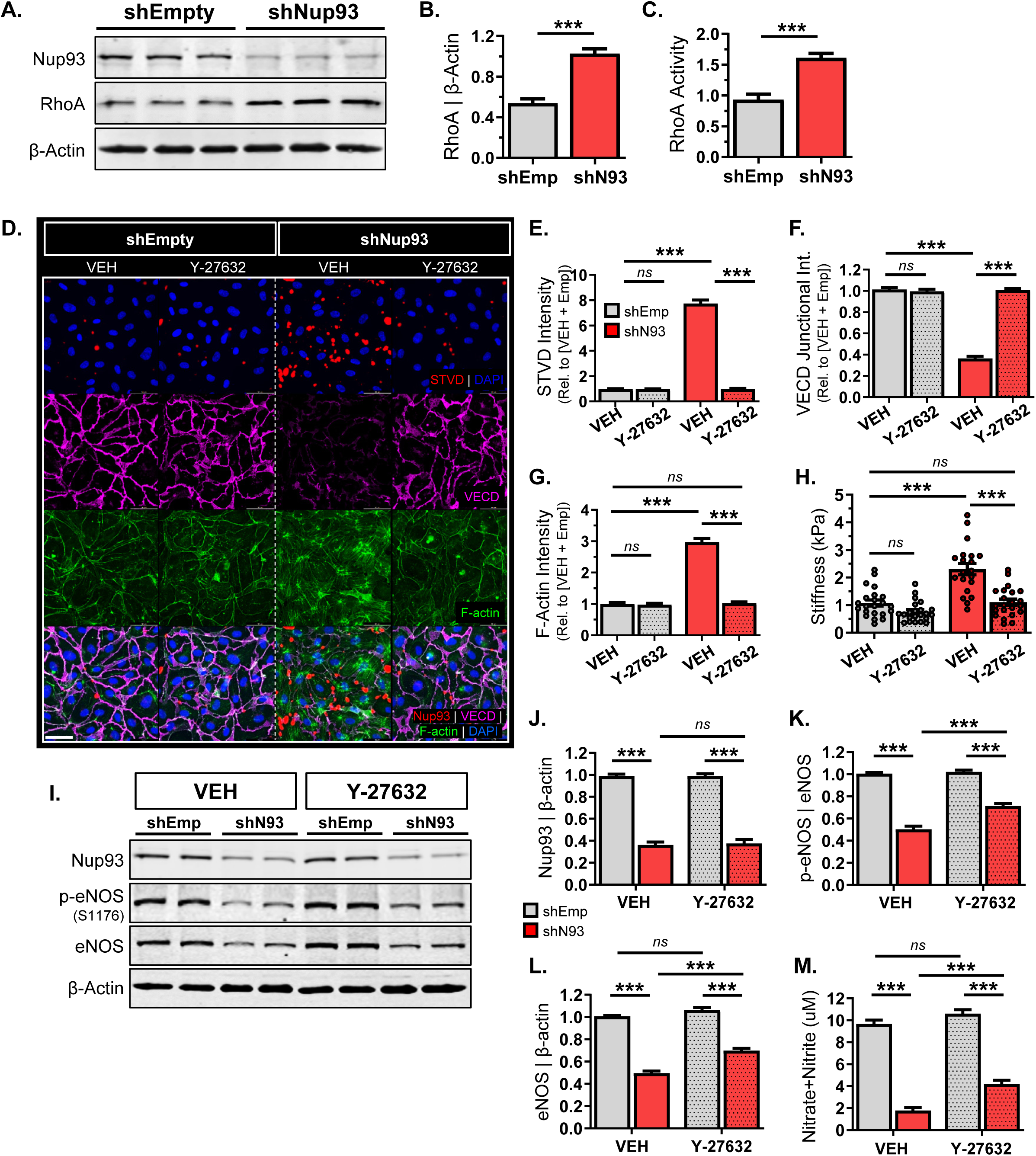
Endothelial Nup93 deficiency leads to RhoA/ROCK hyperactivity. **(A)** Nup93 knockdown significantly increases endothelial RhoA protein levels, as quantified in **(B)**. **(C)** Loss of Nup93 also increases GTP-loaded RhoA levels to indicate heightened RhoA activity. **(D)** shNup93-transduced ECs treated with Y-27632 (10µM; 24hrs) exhibit decreased streptavidin staining, less stress fiber formation, and enhanced VE-cadherin junctional intensity. Quantified in **(E-G)**. (**H)** Y-27632 treatment in Nup93-deficient ECs reduces cellular stiffness, as AFM readouts indicate values similar to that of control conditions. (n=3, 8 reads per group/expt). **(I)** Y-27632 exposure in Nup93-depleted ECs partially rescues both phosphorylated and total eNOS levels, whereas Nup93 remains unaffected (*shown in duplicates*). Quantified in **(J-L)**. **(M)** Conditioned media from Y-27632-treated Nup93-deficient ECs show a partial albeit significant restoration in secreted NO levels. n=3, *\*\*\* p<0.001*

We next evaluated the impact of pharmacological Rho/ROCK inhibition on eNOS expression and function in Nup93-deficient ECs. Similar to previous results (**Fig.3**), Nup93 knockdown significantly reduced the protein levels of both phosphorylated and total eNOS in vehicle-treated conditions (**Fig.5I-K**). Additional treatment with Y-27632 does not affect Nup93 protein; Nup93 levels remain depleted regardless of RhoA/ROCK suppression (**Fig.5I&J**). Pharmacological inhibition of RhoA/ROCK, however, results in a partial, though significant, rescue of both phosphorylated and total eNOS levels when compared to vehicle-treated Nup93-deficient ECs (**Fig.5I,K,L**). Furthermore, Y-27632 exposure in Nup93-deficient ECs results in a partial rescue of eNOS enzymatic function as evidenced by the increased NO levels detected from conditioned media (**Fig.5M**), albeit still less than healthy controls. Nonetheless, these results identify RhoA/ROCK hyperactivation in Nup93-deficient ECs as a contributing factor in eNOS dysregulation, where RhoA/ROCK inhibition is marginally sufficient to restore eNOS function.

Endothelial senescence is characterized by a spectrum of biomarkers and maladaptive functional changes^49,54^. Our recent reports identify Nup93 loss and dysregulated nucleocytoplasmic transport as a major consequence of endothelial aging. As such, targeted knockdown of Nup93 increased the expression of well-accepted senescence biomarkers (*i.e.,* SA-βgal) with corresponding defects in nuclear transport^14^. It is, however, unclear whether the inhibition of RhoA/ROCK activity can suppress biomarkers of senescence (*i.e.,* SA-βgal) and/or restore nucleocytoplasmic transport. We therefore treated Nup93-deficient (and control) ECs with Y-27632 for subsequent detection of SA-βgal. Targeted knockdown of Nup93 significantly increased the percentage of SA-βGal-positive ECs (**Fig.S3A&B**), as previously reported^14^. Following the same Y-27632 treatment strategy, inhibition of RhoA/ROCK did not, however, prevent the SA-βGal expression in Nup93-depleted ECs (**Fig.S3A&B**). We additionally assessed the impact of RhoA/ROCK inhibition on nuclear transport using an artificial steroid-responsive GFP-tagged glucocorticoid receptor (RGG)^29^. In brief, steroid-deficient conditions prevent the exposure of a nuclear localization sequence, therefore resulting in cytoplasmic preference for the GFP reporter. As such, healthy, vehicle-treated ECs exhibit very little nuclear GFP signal to indicate intact nucleocytoplasmic transport properties (**Fig.S3C&E**). Targeted loss of Nup93, however, leads to a significant accumulation in nuclear GFP signal to signify impaired NPC function (**Fig.S3C-E**), as previously reported^14^. Additional treatment with Y-27632 does not restore nucleocytoplasmic transport function; Nup93-deficient ECs continue to exhibit predominantly nuclear-enriched GFP signal despite the presence of a RhoA/ROCK inhibitor (**Fig.S3C-E**). While suppression of RhoA/ROCK activity does not seemingly restore NPC transport function, mitigating RhoA/ROCK activity sufficiently rescues cytoskeletal-reliant behavior. Collectively, these observations identify Nup93 loss as a preceding event that may initiate EC senescence via the consequent perturbation of multiple cellular cascades, with RhoA/ROCK overactivation selectively mediating cytoskeletal-dependent defects.

### Endothelial loss of Nup93 leads to Sun1 decline for enhanced RhoA/ROCK signaling

Thus far, our findings detail an inverse relationship between endothelial Nup93 protein levels and RhoA/ROCK signal activity. While we identify Nup93 as a necessary player in limiting endothelial RhoA signaling, the underlying mechanism remains unclear. Intriguingly, recent studies have begun to implicate nuclear envelope proteins (*i.e.,* LINC proteins) as novel regulators of endothelial function. Critical for nucleocytoskeletal connection, LINC complex proteins (*i.e.,* Sun1) have been shown to associate with nucleoporins^55^ and alter RhoA activity to regulate the actin cytoskeleton^56^. Supporting a role for nuclear envelope proteins in regulation of endothelial behavior, loss of Sun1 was recently reported to trigger EC permeability and stress fiber formation via hyperactivation of the RhoA/ROCK cascade^6^. Given the similar phenotypes (**Fig.3**), we therefore measured Sun expression levels in Nup93-deficient ECs. Alluding to an NPC-LINC communication axis, we find that loss of Nup93 leads to a concomitant decrease in endothelial Sun1 protein, whereas Sun2 remains largely unaffected (**Fig.6A-C**). Moreover, primary MLECs isolated from Nup93iECKO mice also exhibit a prominent reduction in Sun1 with no effect on Sun2 protein levels (**Fig.6D-F**), corroborating *in vitro* results. Intriguingly, both Sun1 and Sun2 mRNA levels, nonetheless, remain unchanged with Nup93 knockdown (**Fig.S4A-C**) to indicate Sun1 loss via a post-transcriptional mechanism.

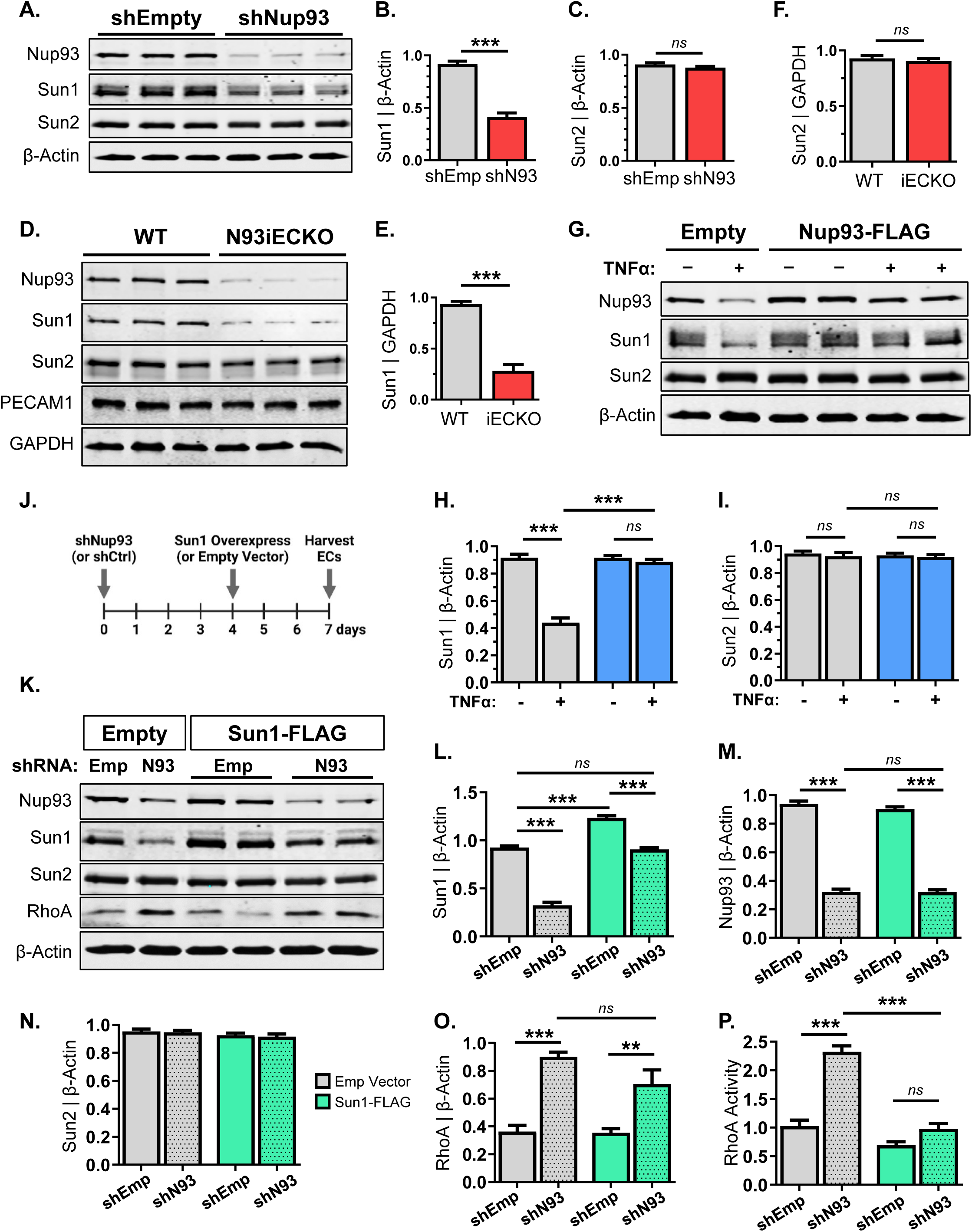

Our recent reports identify an age-associated decline in endothelial Nup93 protein, adding vascular cells to the growing list of cell types that exhibit nuclear permeability with age^12,14^. While our studies attributed impaired nucleocytoplasmic transport as the major mechanism of endothelial dysfunction, it remains unclear whether an NPC-LINC communication axis is also perturbed with EC senescence. We therefore revisited an *in vitro* senescence model^48^ and again observed a visible decline in Nup93 protein levels (**Fig.6G**), similar to our previous reports^14^. We find that endothelial exposure to chronic inflammation also led to a selective decrease in Sun1 protein levels (**Fig.6G-I**). Resembling a possible dependency on Nup93 protein, we next revisited our Nup93 re-expression model (**Fig.4E**) to address whether Nup93 restoration could also rescue Sun1. Intriguingly, restoring Nup93 protein to baseline levels in already senescent ECs led to a full recovery of Sun1 protein; Sun1 levels in TNFα-treated ECs are comparable to that of healthy controls upon exogenous Nup93 delivery. Sun2, nevertheless, remains unaffected regardless of the condition (**Fig.6G-I**).

To establish a unidirectional cascade and definitively establish Sun1 as a downstream player of Nup93, similar lentiviral-based methods were used to restore Sun1 levels in Nup93-deficient ECs. In brief, ECs were first infected with shRNA lentiviral constructs (shEmpty & shNup93), followed by exogenous lentiviral infection of Sun1 (Empty & Sun1-FLAG), as shown in **Figure 6J**. Lentiviral-mediated expression of the Sun1-FLAG construct results in proper localization (**Fig.S4D**) and a significant increase, albeit only a 1.3-fold overexpression, when compared to endogenous levels. More importantly, Sun1-FLAG delivery in Nup93-deficient ECs fully restores Sun1 to endogenous levels; Sun1 expression is comparable to that of heathy EC controls (**Fig.6K&L**). Nup93 levels, however, remain significantly decreased regardless of exogenous Sun1 expression (**Fig.6K&M**). Furthermore, restoring Sun1 levels does not mitigate RhoA protein expression, as RhoA expression remains elevated (**Fig.6K&O**). Nonetheless, we find that restoring Sun1 mitigates RhoA activity levels in Nup93-deficient ECs (**Fig.6P**) to suggest Nup93-Sun1 may additionally engage alternative RhoA regulatory mechanisms. Sun2 levels continue to remain unaltered across all conditions (**Fig.6K&N**).

### Restoring Sun1 levels in Nup93-deficient ECs rescues EC barrier integrity and eNOS function

Endothelial permeability is regulated in part by the cellular cytoskeleton and contractility. Previous work has shown that decreasing endothelial contractility restores monolayer integrity. Implicated in EC contraction, inhibition of aberrant RhoA/ROCK activity can mitigate age-associated vascular stiffening and improve barrier function in aged mice^57^. Nup93-deficient ECs indeed show an increase in monolayer permeability, and complete rescue upon inhibition of RhoA/ROCK activity (**Fig. 5D**). While these studies identify RhoA/ROCK hyperactivation in Nup93-depleted ECs as the driver of barrier disruption, the regulation of EC barrier function via an NPC-LINC axis remains elusive. We therefore re-expressed Sun1, a repressor of RhoA/ROCK signaling, in Nup93-deficient ECs (**Fig.6J**) for subsequent permeability studies. Similar to previous results (**Fig.5**), we find that targeted loss of Nup93 significantly enhances paracellular permeability, as indicated by the visible increase in streptavidin staining and decreased VE-cadherin junctional localization (**Fig.7A-C**). Nup93 loss again results in increased stress fibers and endothelial stiffening (**Fig.7D&E**). Interestingly, restoring Sun1 levels significantly reduced streptavidin signal and fully restored junctional VE-cadherin expression in Nup93-deficient ECs, similar to that of healthy control ECs (**Fig.7A-C**). Exogenous Sun1 delivery also attenuated stress fiber formation and reduced cellular stiffness, as EC rigidity in Nup93-deficient ECs is indistinguishable from healthy controls ECs (**Fig.7A,D,E**). Hence, RhoA/ROCK hyperactivation and consequent monolayer permeability in Nup93-depleted ECs likely reflects the associative decline in Sun1 protein, thereby indicating a Nup93-Sun1 communication axis for whole-cell regulation.

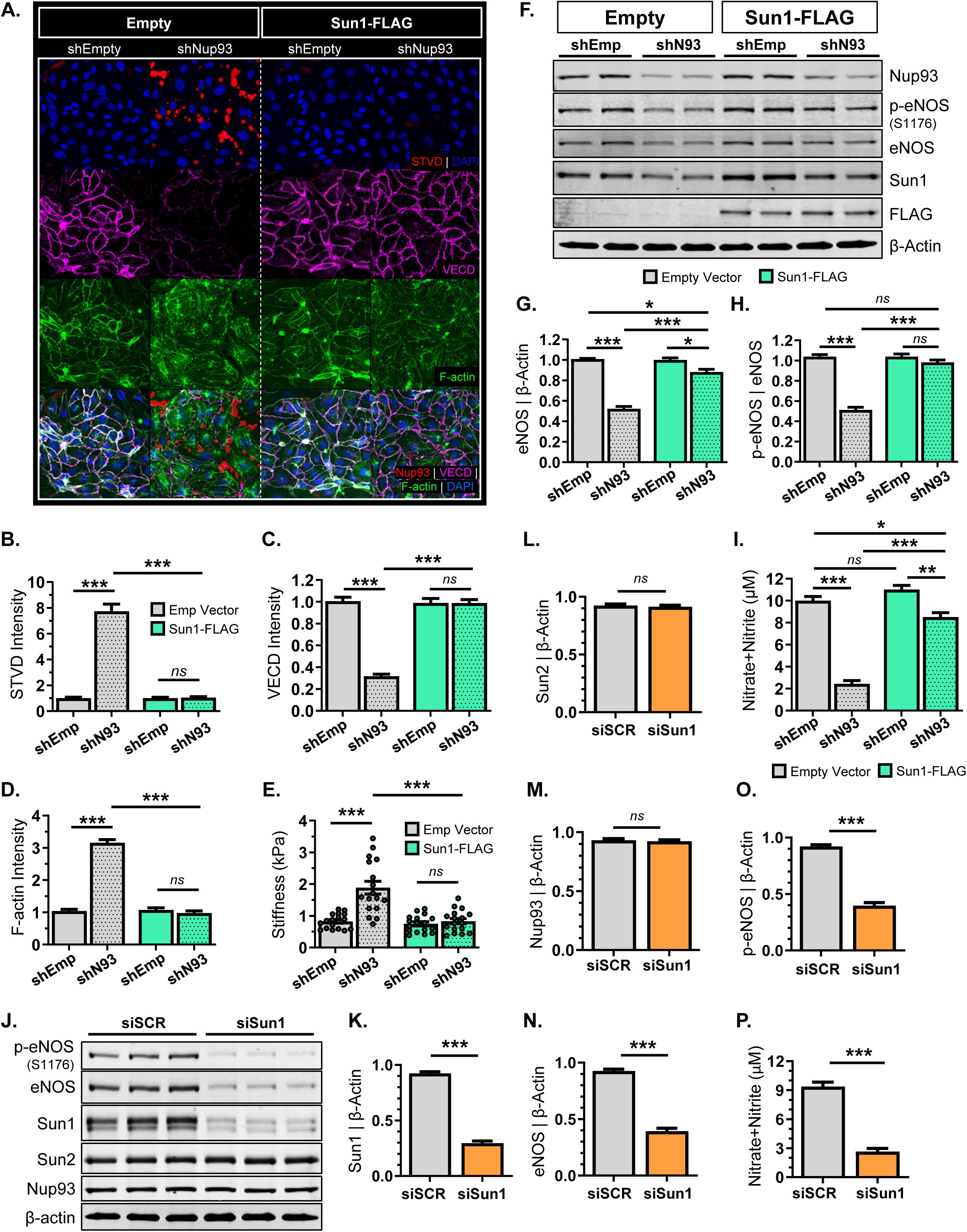

Using the Sun1 restoration model, we next sought to investigate the NPC-LINC axis in nucleocytoplasmic transport function and EC senescence. As previously reported^14^ and shown herein (**Fig.S3A**), targeted knockdown of Nup93 significantly increases the percentage of SA-βGal positive cells to confirm EC senescence. Re-expression of Sun1, however, does not influence SA-βGal expression, as levels remain high regardless of Sun1 restoration (**Fig.S5A&B**). To examine the NPC-LINC axis on nucleocytoplasmic transport, primary ECs were again transduced with the GFP-tagged nucleocytoplasmic RGG reporter construct. An *in vitro* senescence model together with Sun1 restoration was utilized, similar to the rescue schematic in **Figure 4E**. In brief, RGG-expressing ECs were exposed to long-term inflammation (TNFα [10ng/mL]; 6 days) to induce senescence, followed by exogenous Sun1-FLAG infection. Indicative of proper nucleocytoplasmic transport function, healthy EC controls (in the absence of steroids) show a predominant cytoplasmic localization of GFP signal. Aligning with previous reports^14^, chronic TNFα treatment, however, significantly decreases Nup93 and Sun1 expression, resulting in nuclear localization of GFP signal to indicate impaired transport function. Intriguingly, exogenous delivery of Sun1 protein in already senescent ECs, as confirmed by immunofluorescent staining, does not restore NPC function; GFP signal remains still predominantly in the nucleus (**Fig.S5C-E**). Taken together, restoring Sun1 in Nup93-deficient ECs cannot rescue endothelial senescence nor select cellular functions, namely nucleocytoplasmic transport. These results insinuate loss of Nup93 as the initiating driver of EC dysfunction, where increases in SA-βGal expression and disrupted NPC transport may occur independent of Sun1.

Using both *in vitro* and *in vivo* models, we find that loss of endothelial Nup93 significantly decreases eNOS levels. Notably, endothelial-targeted Nup93 knockdown mice also exhibit decreased circulating NO levels and impaired NO-dependent vasodilatory responses. While several studies implicate nuclear envelope proteins in endothelial behavior, the role of nuclear envelope components in whole vessel physiology remains undefined. We therefore utilized the Sun1 restoration model (**Fig.6J**) to investigate the NPC-LINC axis in eNOS regulation. Loss of Nup93 drastically diminishes both phosphorylated and total eNOS, as previously shown (**Fig.3F**, **Fig.7F-H**). Intriguingly, restoring Sun1 in Nup93-deficient ECs leads to a partial rescue in total eNOS, whereas eNOS phosphorylation appears fully restored (**Fig.7F-H**). Moreover, exogenous Sun1 expression significantly increased NO production in Nup93-deficient ECs; NO levels in conditioned media near that of healthy controls ECs (**Fig.7I**). To directly assess the role of Sun1 in eNOS regulation, siRNA targeted strategies were used. Immunoblotting validates a selective loss in Sun1 protein, as Sun2 protein levels remain unchanged (**Fig.7J-L**). Interestingly, Nup93 also remains unaffected with loss of Sun1 (**Fig.7M**). We find, however, that Sun1 knockdown significantly decreased total and phosphorylated eNOS levels (**Fig.7J,N,O**). We additionally collected conditioned media prior to protein harvesting, where results show a corresponding decrease in NO production in Sun1 knockdown ECs (**Fig.7P**). This data identifies Sun1 loss – a consequence of Nup93 depletion – as a mechanism of eNOS impairment to implicate nuclear envelope proteins in eNOS and NO regulation. Hence, loss of Nup93, a natural consequence of endothelial aging, disrupts a range of key cellular functions to accumulate a cascade of dysregulated endothelial behaviors. We herein uncover the existence of an NPC-LINC axis, necessary for nucleus-to-periphery molecular communication and EC health, that is inadvertently disrupted with endothelial Nup93 loss.

## DISCUSSION

As the innermost lining of the vasculature, ECs are constantly subjected to systemic inflammation and particularly vulnerable to aging^58,59^. Expanding upon our recent findings, this study provides incremental evidence supporting Nup93 as a major regulator of EC health. We herein identify several novel findings to support the premise of a Nup93-Sun1 communication axis as a novel mechanism coupling endothelial NPC biology to nucleocytoskeletal regulation. First, using both *in vitro* and *in vivo* approaches, we find that loss of endothelial Nup93 leads to RhoA/ROCK hyperactivity and consequent disruption of endothelial barrier function for vessel leakage. Second, we provide evidence that a decline in endothelial Nup93, whether using chronic inflammation or targeted knockdown methods, results in a corresponding decrease in eNOS protein, thus blunting NO bioavailability and vessel vasodilatory responsiveness. These observations suggest a mechanism of eNOS dysregulation and EC dysfunction that emerges with endothelial Nup93 loss. Lastly, we herein identify loss of Sun1, a major component of the LINC complex and a known inhibitor of RhoA/ROCK signaling^60^, as an outcome of Nup93 decline that contributes to eNOS reduction. While the precise mechanisms underlying Nup93 and Sun1 protein decline have not yet been delineated, our findings identify a mechanism of eNOS dysregulation prompted by Nup93 deficiency. Overall, these studies highlight the existence of an NPC-LINC axis, necessary for nucleus-to-periphery molecular communication and EC health, that is inadvertently disrupted with Nup93 loss.

Senescence of vascular ECs is accompanied by perturbations in cytoskeletal integrity, resulting in endothelial permeability and increased stiffening^49^. We herein demonstrate impaired endothelial barrier function and increased EC rigidity with loss of Nup93, confirming the manifestation of age-associated cytoskeletal defects. RhoA/ROCK hyperactivity is a major contributor to age-associated vasoconstriction and hypertension^57,61^. We establish RhoA/ROCK hyperactivity and endothelial stiffening as a consequence of Nup93 loss, akin to previous reports of endothelial aging. We find that restoring Sun1 in Nup93-deficient ECs rescued various cytoskeletal-dependent EC behaviors, including the proper assembly of F-actin. Underscoring a role for Nup93 endothelial actin dynamics, these observations implicate Nup93, through Sun1 regulation, in RhoA/ROCK-mediated cytoskeletal remodeling. Exogenous Sun1 expression, however, does not impact RhoA protein levels. Rather, re-expression of Sun1 reduced RhoA enzymatic activity despite elevated RhoA protein levels. Mechanistically, endothelial Sun1 was reported to stabilize microtubules, thereby sequestering RhoGEF-H1 - a known RhoA effector - for inactivity^60^. Hence, restoring Sun1 in Nup93-deficient ECs likely re-establishes the EC cytoskeletal architecture to properly sequester RhoGEF from the cytoplasm. This would maintain RhoA in an inactive form despite the elevated RhoA protein levels, emphasizing the importance of protein enzymatic activity rather than total levels. Moreover, our studies begin to illustrate Sun1 as a downstream effector of Nup93, where loss of Nup93 initiates Sun1 degradation for RhoA/ROCK hyperactivity.

Nuclear envelope proteins have gained traction as novel regulators of endothelial health to include our recent reports identifying Nup93 loss as a natural outcome of endothelial aging. Given its canonical role in maintaining NPC structure, nucleocytoplasmic transport defects stands as the major consequence of Nup93 loss in various cell types, including ECs^14,62^. Recent investigations, however, indicate nucleoporins can also regulate transport-independent features to include cellular mechanobiology^63^. In other words, age-associated loss of Nup93 may initiate tiers of cellular dysregulation, with dominant consequences on nucleocytoplasmic transport, as detailed in our recent reports^14^, and secondary effects on nucleocytoskeletal integrity as we show herein. In support of this premise, we find that nucleocytoplasmic transport remains significantly impaired in Nup93-deficient ECs, regardless of Sun1 restoration. Barrier leakage and endothelial stiffness incurred with Nup93 deficiency is, however, fully reversed when using either pharmacological RhoA/ROCK inhibition or Sun1 protein restoration methods. We find that either method is sufficient, and effective, to fully rescue cytoskeletal-dependent properties in Nup93-deficient ECs. In contrast, eNOS protein levels and NO bioavailability remain only partially recovered when using either rescue strategy. It is indeed surprising to observe complete rescue in select endothelial paradigms, despite striking defects in nucleocytoplasmic shuttling. Nonetheless, these observations begin to reveal the transport-independent features of Nup93. Intriguingly, cytoskeletal remodeling seemingly stands as the underlying commonality in the fully rescued endothelial paradigms with Sun1 restoration. As such, other vital endothelial functions (*i.e.,* NO production) are only partially rescued to suggest disruption of alternative eNOS regulatory mechanisms reliant on NPC function. Loss of Nup93 may effectively trigger eNOS disruption through multiple modalities, engaging both cytoskeletal-mediated mechanisms (*i.e.,* RhoA/ROCK activation) and impairing NPC shuttling of eNOS transcriptional regulators (*i.e.,* KLF2/4, MRTF)^64,65^. Nonetheless, restoring Sun1 protein had a greater rescue effect than RhoA/ROCK inhibition to support Sun1, and proper nucleocytoskeletal architecture, as a more efficient upstream modulator of RhoA/ROCK activity. Furthermore, these studies establish Sun1 decline, a collateral event of Nup93 loss, as a disruptor of eNOS function that occurs independent of Nup93-mediated NPC transport function. Future studies will nonetheless be necessary to fully understand how chronic inflammation tampers the Nup93-Sun1 axis.

Lastly, it is important to recognize that the nuclear envelope not only functions for compartmentalization, but also as a highly dynamic force sensor^3^. Involving a range of transcriptional and posttranslational mechanisms, laminar shear flow remains one of the most important determinants for continuous NO generation conferring EC and vascular protection^66^. Endothelial actin cytoskeletal remodeling occurs upon mechanical stimulation^67^, where F-actin integrity is required for laminar blood flow-induced NO production^68^. Our results herein identify stress fiber formation and increased endothelial stiffening with endothelial loss of Nup93, substantiating recent studies implicating Nup93 in cytoskeletal actin regulation^69^. While proper Nup93 expression is without a doubt essential for EC health, it remains virtually unknown how endothelial Nup93 dysregulation precipitates vascular disease pathogenesis. Collectively, these prior studies, together with our results herein, implicate a role for Nup93 in endothelial flow adaptation, where suboptimal levels of endothelial Nup93 may blunt the benefits of laminar flow to impair both cellular alignment and nucleocytoplasmic transport. As such, our findings insinuate endothelial Nup93 loss, and consequent eNOS disruption, as a trigger for systemic hypertension and atherosclerotic disease progression, a topic for future reports. Furthermore, our *in vitro* studies identify Nup93 restoration as a plausible method to rescue key EC behaviors. It remains, nevertheless, unclear whether exogenous Nup93 expression could mitigate vascular disease. Further investigations will hence be necessary to fully understand the multiscale impact and therapeutic potential of Nup93 delivery in reestablishing vascular health.

In conclusion, we identify RhoA/ROCK-mediated eNOS disruption as a repercussion of Nup93 to highlight the existence of a regulatory mechanism between NPC components and eNOS function. We report that Nup93 loss triggers a concomitant decline in Sun1 levels, thereby driving RhoA/ROCK hyperactivity for consequent stress fiber formation, cellular stiffening, and endothelial leakage. Demonstrating the unidirectionality of Nup93-Sun1 regulation, we show that exogenous Sun1 delivery in Nup93-deficient ECs, despite impaired NPC transport, fully reverses cytoskeletal-related endothelial defects. These findings not only suggest that the restoration of the LINC complex and proper nucleocytoskeletal cellular architecture but begin to shed light on Nup93-mediated NPC transport-independent mechanisms. Our findings signify, for the first time, a mechanism for eNOS dysregulation prompted by Nup93 deficiency. In conclusion, we demonstrate a critical connection between nuclear envelope perturbations and eNOS disruption to indicate the presence of an eNOS feedback mechanism that originates at the NPC.

## AFFILIATIONS

Department of Physiology and Biophysics, University of Illinois Chicago (T.D.N., Y.Z.K, R.M., J.M. B, J.M., M.A.W., M.Y.L.). Center for Cardiovascular Research, University of Illinois Chicago (T.D.N, J.M., S.A.P., M.Y.L.). Department of Biomedical Engineering, University of Illinois Chicago (F.H., J.C.L.).

## ACKNOWLEDGEMENTS

We thank Patrick Lusk, Ph.D. (Yale University) for his invaluable advice on NPC biology.

## SOURCES OF FUNDING

This work was supported in part by the National Institutes of Health (R00 HL130581 to M.Y.L.), the American Heart Association (AHA 941057 to M.Y.L.)., and from UIC (AGR/Diess to T.D.N.).

## DISCLOSURES

None.

## AUTHOR CONTRIBUTIONS

T.D.N performed and analyzed most of the data presented herein. Y.Z.K. performed the NO assays. F.H. and J.C.L. performed and consulted on the AFM studies, respectively. R.M. performed the RT-qPCR experiments. J.M.B. performed the SA-βGal assays. J.M. harvested and sectioned mouse tissues for immunostaining. M.A.W generated constructs and reagents. L.M. and S.A.P. performed and consulted on the flow dilation studies, respectively. M.Y.L conceived and directed the project. T.D.N and M.Y.L wrote the manuscript, which was reviewed and edited by all authors.

## Notes

### Competing Interest Statement

The authors have declared no competing interest.

