## Supplementary material for "Endothelial Nucleoporin93 (Nup93) Maintains Vascular Function via Sun1-Dependent Regulation of RhoA-eNOS Signaling": Supp Fig/Tables

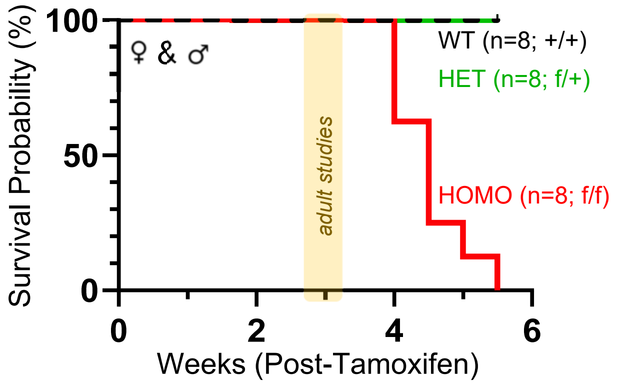


**Figure S1. Endothelial loss of Nup93 is incompatible with life.** Mice homozygous for the Nup93 floxed allele invariably face lethality irrespective of sex whereas heterozygous Nup93 floxed mice remain grossly unaffected. All *in vivo* adult studies were performed within 3 weeks of the last tamoxifen injection to avoid the morality time window (denoted in yellow).

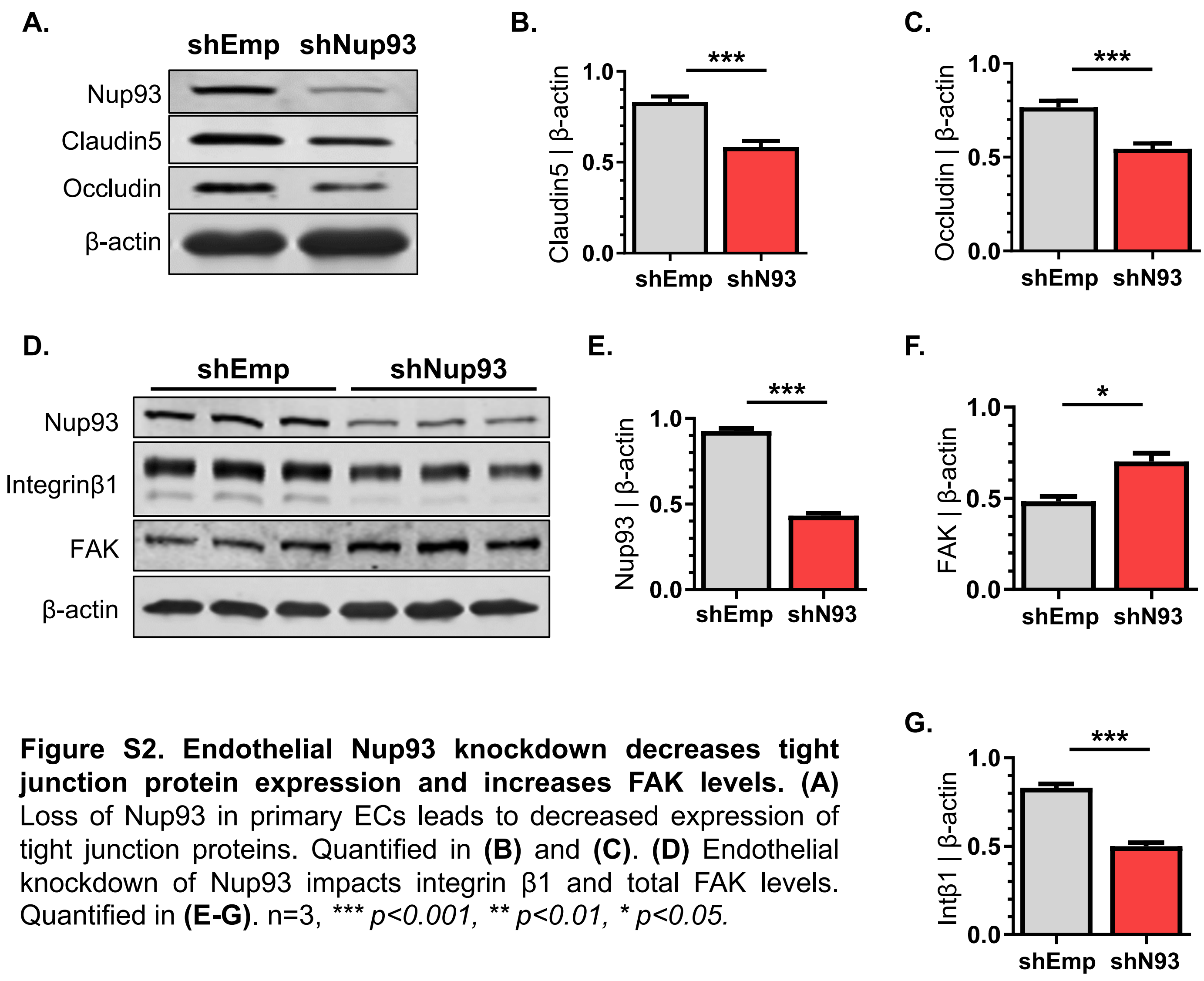


**Figure S2. Endothelial Nup93 knockdown decreases tight junction protein expression and increases FAK levels. (A)** Loss of Nup93 in primary ECs leads to decreased expression of tight junction proteins. Quantified in **(B)** and **(C)**. **(D)** Endothelial knockdown of Nup93 impacts integrin β1 and total FAK levels. Quantified in **(E-G)**. n=3, *** *p<0.001, ** p<0.01, * p<0.05.*

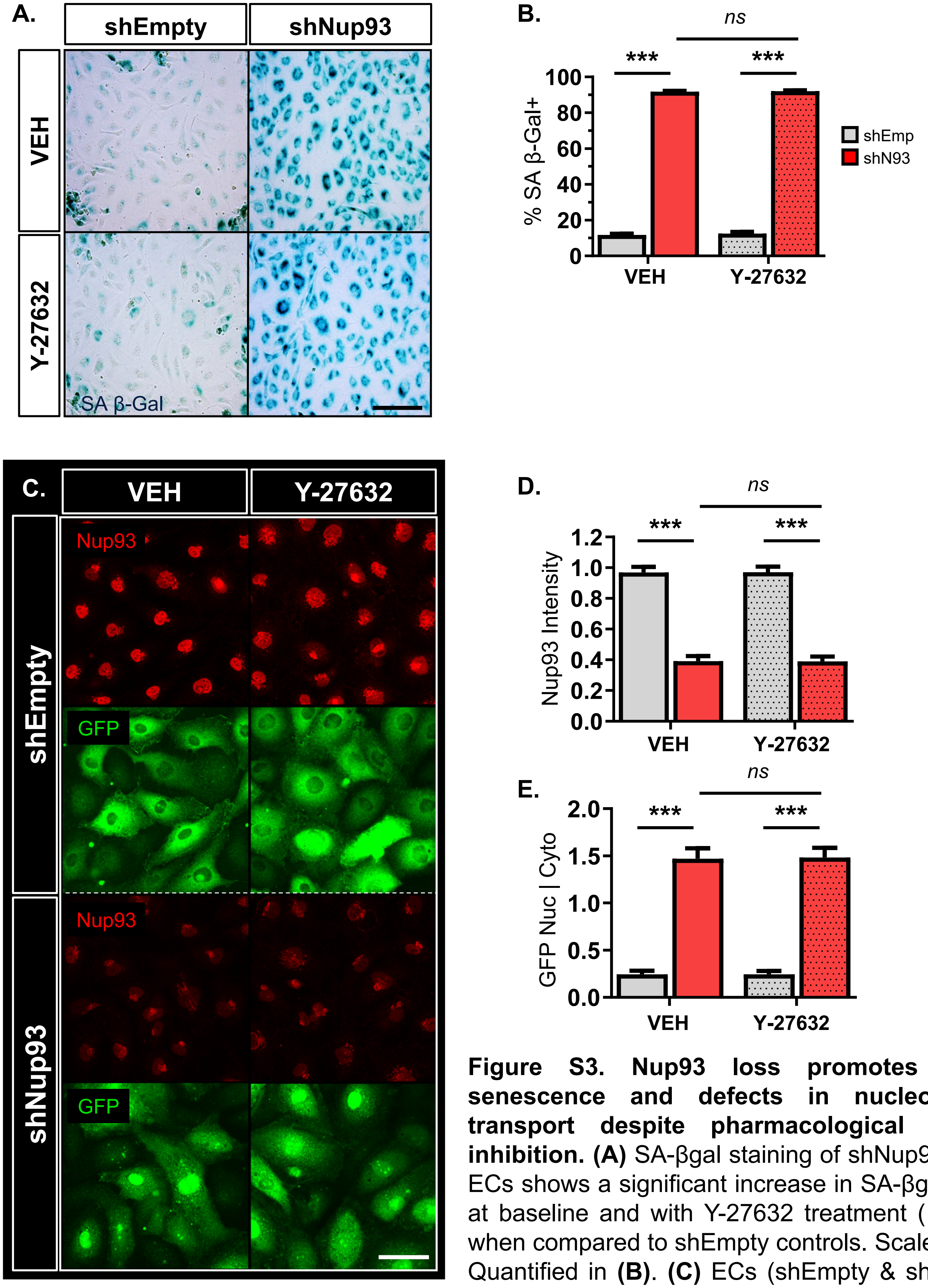


**Figure S3. Nup93 loss promotes endothelial senescence and defects in nucleocytoplasmic transport despite pharmacological RhoA/ROCK inhibition. (A)** SA-βgal staining of shNup93-transduced ECs shows a significant increase in SA-βgal signal both at baseline and with Y-27632 treatment (10μM; 24hrs) when compared to shEmpty controls. Scale bar=200μm. Quantified in **(B)**. **(C)** ECs (shEmpty & shNup93) were further transduced with the RGG construct using lentiviral methods followed by Y-27632 treatment (10μM, 24hrs). GFP nuclear-to-cytoplasmic intensity remains unaffected by Y-27632 treatment in Nup93 knockdown ECs when compared to vehicle-treated conditions. Scale bar=50μm. Nup93 expression and GFP nuclear-to-cytoplasmic levels quantified in **(D)** and **(E)**. n=3, *** *p<0.001*

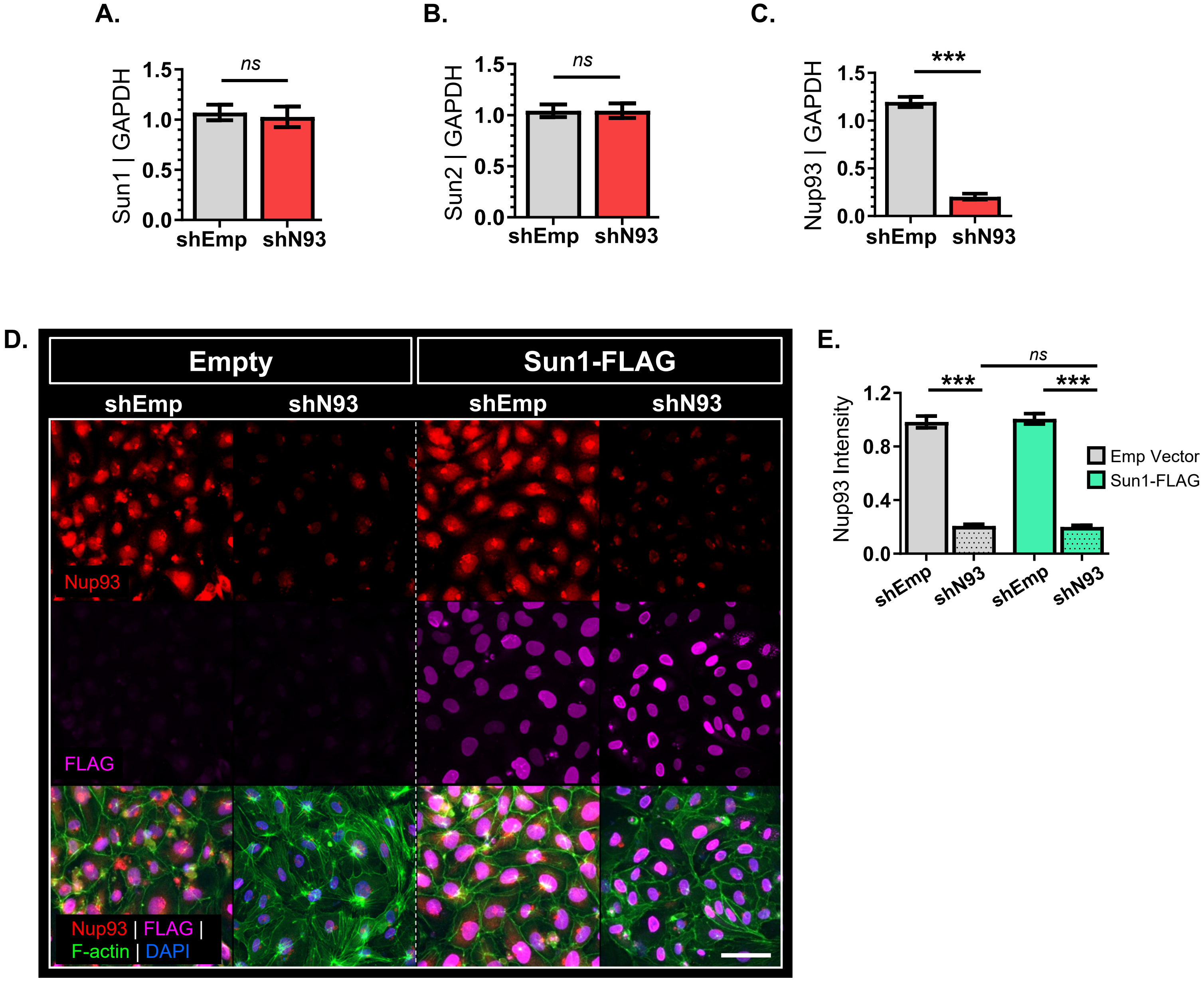


**Figure S4. Targeted knockdown of endothelial Nup93 does not affect Sun1/2 transcript levels. (A-C**) RT-qPCR analysis in primary ECs indicates a significant decrease in Nup93 mRNA levels when using shRNA-based method, whereas Sun1 and Sun2 expressions remain unaffected. **(D)** Immunofluorescence staining validates shRNA-mediated Nup93 loss and proper localization of exogenous Sun1 in both shEmpty control and shNup93 ECs, as demonstrated via nuclear FLAG signal. Nup93 signal quantified in **(E)**. Scale bar=50μm. n=3, *** *p<0.001*

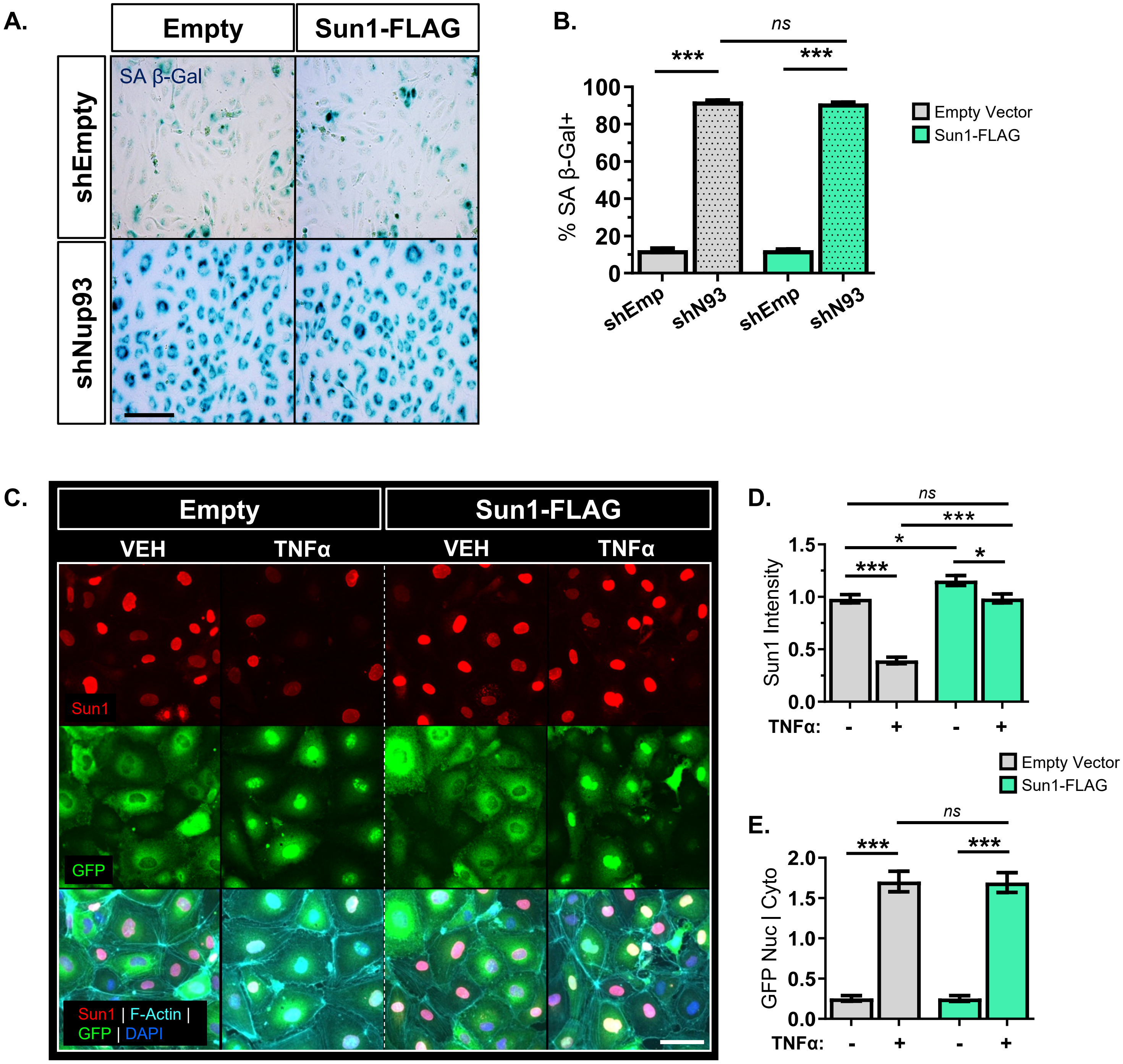


**Figure S5. Nup93 loss triggers endothelial senescence and defects in nucleocytoplasmic transport despite restoration of Sun1 levels. (A)** SA-βgal staining of shNup93-tranduced ECs shows a significant increase in SA-βgal signal regardless of exogenous Sun1 delivery. Scale bar=200µm. Quantified in **(B)**. **(C)** RGG-transduced primary ECs were first exposed to chronic inflammation (TNFα [10ng/mL]; 6 days) to induce EC senescence and nucleocytoplasmic transport defects. While long-term inflammation significantly reduces Sun1 expression, restoring Sun1 does not rescue NPC transport function; GFP nuclear-to-cytoplasmic intensity remains unaffected with exogenous Sun1 expression. Quantified in **(D)** and **(E).** Scale bar=50µm. n=3, *** *p<0.001, * p<0.05*

**Table S1. Genotyping primers**

| Target Gene | Forward (5'-3') | Reverse (5'-3') | Product Size (bps) |
|---|---|---|---|
| ***Mouse*** | | | |
| ***Nup93 flox*** | TTTGGGTACCTTACTCCCACA | CAACCCAATGAGTCCCTTCT | 552 (WT); 620 (flox) |
| ***Generic Cre*** *(JAX, oIMR1084 & oIMR1085)* | GCGGTCTGGCAGTAAAAACTATC | GTGAAACAGCATTGCTGTCACTT | 100 |
| ***Internal Control*** *(JAX, oIMR7338 & oIMR7339)* | CTAGGCCACAGAATTGAAAGATCT | GTAGGTGGAAATTCTAGCATCATCC | 324 |

**Table S2. RT-qPCR primers**

| Target Gene | Forward (5'-3') | Reverse (5'-3') | Product Size (bps) |
|---|---|---|---|
| ***Human*** | | | |
| *NUP93* | AGGACAATGCCCTGCTGTCT | AAGGGCGTCTTCTCCTGATG | 152 |
| *SUN1* | GGACGTGTTTAAACCCACGACTTCTCG | CTCTGACTTTAGCTGATCCAGCTCCAGC | 457 |
| *SUN2* | AAACTGCTGCTCGCATCC | GAGTCTTGCTGATGCTCTGCT | 81 |
| *GAPDH* | CTCTCTGCTCCTCCTGTTCGAC | TGAGCGATGTGGCTCGGCT | 71 |
